# Leukocyte-type 12/15-lipoxygenase is essential for timely inflammation-resolution and effective tissue regeneration following skeletal muscle injury

**DOI:** 10.1101/2025.05.13.653766

**Authors:** Binayok Sharma, Xinyue Lu, Hamood Rehman, Vandré C. Figueiredo, Carol Davis, Holly Van Remmen, Shihuan Kuang, Susan V. Brooks, Krishna Rao Maddipati, James F. Markworth

**Author notes:** Contributed equally (Binayok Sharma, Xinyue Lu, Hamood Rehman). Co-senior authors (Krishna Rao Maddipati, James F. Markworth). **Corresponding authors:** James F. Markworth, Ph. D, Krishna Rao Maddipati, Ph.D.

## Abstract

Unlike traditional anti-inflammatory therapies which may interfere with musculoskeletal tissue repair, pharmacological administration of specialized pro-resolving lipid mediators (SPMs) can promote timely resolution of inflammation while stimulating skeletal muscle regeneration. Despite this, the potential role of endogenous inflammation-resolution circuits in skeletal muscle injury and repair remains unknown. Here, we investigated the effect of whole-body knockout of leukocyte-type 12/15-lipoxygenase (12/15-LOX) on acute inflammation and regeneration following skeletal muscle injury in mice. Prior to muscle injury, *Alox15*^-/-^ mice displayed lower intramuscular concentrations of 12/15-LOX-derived lipid mediators than wild type (WT) mice, and this was associated with chronic low-grade muscle inflammation. *Alox15*^-/-^ mice mounted an exaggerated acute immune response to sterile skeletal muscle injury which was associated with a local imbalance of pro-inflammatory vs. pro-resolving lipid mediators. During the regenerative phase, *Alox15^-/-^* mice displayed defects in myogenic gene expression, myofiber size, and myonuclear accretion. Mechanistically, bone marrow-derived macrophages (MФ) obtained from *Alox15*^-/-^ mice produced less 12/15-LOX-derived lipid mediators and this was associated with impaired M2 polarization. Isolated myogenic progenitor cells also produced many LOX metabolites in response to long chain polyunsaturated fatty acid (LC-PUFA) supplementation, including bioactive SPMs. *Alox15*^-/-^ myoblasts were both impaired in their ability to produce SPMs and were insensitive to the stimulatory effect of LC-PUFAs on *in vitro* myogenesis. These data show that the 12/15-LOX pathway is essential for timely resolution of acute inflammation and direct determination of myogenic progenitor cell fate following skeletal muscle injury.

## Introduction

Skeletal muscle injury is well-known to induce a robust acute inflammatory response which results in the local appearance of substantial numbers of blood leukocytes^1^. Polymorphonuclear cells (PMNs) (e.g., neutrophils) are among the first blood immune cells to arrive at the site of skeletal muscle injury, usually within minutes to hours^2,3^. The next major type of blood cells that migrate to the site of injury are monocytes that differentiate locally to become tissue macrophages (MФs) which play a crucial role in supporting subsequent myofiber regeneration via immune-muscle cell cross-talk^4–6^.

Lipoxygenase (LOX) enzymes are pivotal in modulating inflammation by oxidizing long chain polyunsaturated fatty acids (LC-PUFAs) to form bioactive lipid mediators^7^. In mammals, specific enzymes are responsible for catalyzing enzymatic oxidation of three sites on arachidonic acid (ARA, 20:4n-6), and thus are named 5-, 12-, and 15-LOX, respectively^8^. 15-LOX oxygenates ARA substrate at carbon-15 to form 15-hydroperoxy-eicosatetranoic acid (15-HpETE), which is further reduced to 15-hydroxy-eicosatetranoic acid (15-HETE)^9^. However, certain LOX enzymes possess dual specificities, resulting in differences among species^8^. Importantly, while the human protein encoded by the *ALOX15* gene (15-LOX-1) produces mainly 15-HETE, the analogous protein encoded by the murine *Alox15* gene produces both 12-HETE and 15-HETE and is thus often termed 12/15-LOX.

LOX-derived lipid mediators are involved in a myriad of physiological and pathological processes, including the initiation and resolution of the inflammatory response^10^. Monohydroxylated PUFA metabolites such as 5-, 12-, and 15-HETE may exert direct biological effects^11^. In addition, certain monohydroxy-PUFAs are potential precursors in the subsequent downstream biosynthesis of a complex array of potential downstream di- and tri-hydroxylated PUFA metabolites^12^. For example, 15-HETE formed via 15-LOX metabolism of ARA can be further metabolized via the 5-LOX pathway to form the lipoxin family of lipid mediators (e.g., LXA_4_)^13^. Unlike classical eicosanoids (e.g., prostaglandins and leukotrienes), the lipoxins have potent anti-inflammatory actions such as limiting PMN chemotaxis and degranulation^14^. The lipoxins are also key molecules acting in the resolution phase of the acute inflammatory response by promoting monocyte recruitment to the site of inflammation^15^ and stimulating MФ mediated phagocytosis of apoptotic PMNs (efferocytosis)^16^.

LC omega-3 (n-3) PUFAs such as eicosapentaenoic acid (EPA, 20:5n-3), docosapentaenoic acid (DPA, 22:5n-3) and docosahexaenoic acid (DHA, 22:6n-3) are also potential substrates for mammalian LOX enzymes. 15-LOX activity converts DHA to 17-hydroxy-docosahexaenoic acid (17-HDoHE)^17^. Subsequently 5-LOX expressing cells (e.g., PMNs) may transform 17-HDoHE to downstream products such as the D-series resolvins (e.g., RvD1)^18^. Similarly, enzymatic metabolism of n-3 DHA by 12-LOX produces 14-hydroxy-docosahexaenoic acid (14-HDoHE) which can be further converted to the maresins (e.g., MaR1)^19^. Based on their key roles in actively bringing about the resolution phase of the acute inflammatory response the lipoxins, resolvins, protectins, and maresins have collectively been termed specialized pro-resolving lipid mediators (SPMs)^20^.

Recent evidence demonstrated by us and independent groups have used liquid chromatography tandem mass spectrometry (LC-MS/MS) profiling to show that many potential LOX pathway metabolites increase locally following acute skeletal muscle injury^21–26^. These novel findings are consistent with earlier reports showing that exercise-induced skeletal muscle damage also transiently increased many LOX metabolites in human blood^27,28^. Despite their clear association with muscle injury and repair processes, the precise role of such LOX-derived lipid mediators in adaptive skeletal muscle remodeling remains uncertain. In the only published study employing a loss-of-function model to date, leukocyte type 12/15-LOX knockout (*Alox15*^-/-^) mice were found to be protected against skeletal muscle wasting induced by denervation surgery^29^. Nevertheless, no prior study has tested the potential impact of loss of 12/15-LOX activity on acute skeletal muscle injury and ensuing myofiber regeneration.

In the current study we investigated the functional role of the *Alox15* gene, which encodes the murine leukocyte-type 12/15-LOX enzyme, on acute skeletal muscle inflammation and regeneration in mouse and cell models. We hypothesized that 12/15-LOX-mediated enzymatic conversion of LC-PUFAs to form bioactive lipid mediator metabolites with anti-inflammatory, pro-resolving, and tissue reparative actions would be essential for timely resolution of the acute innate immune response and robust skeletal muscle regeneration. Our data reveals not only a crucial role of *Alox15* in the successful resolution of acute skeletal muscle inflammation following sterile tissue damage, but also an unexpectedly direct and indispensable role of 12/15-LOX enzyme activity as a key positive determinant of resident myogenic progenitor cell fate within injured muscle.

## Materials and Methods

### Animals

Wild type (WT) C57BL/6J (Jackson, 000664) and whole-body 12/15-LOX knockout (*Alox15*^-/-^) mice on a C57BL/6J background (B6.129S2-*Alox15*^tm1Fun^/J (Jackson, 002778^30^) were sourced from the Jackson Laboratory. Male 6–8-month-old *Alox15*^-/-^ mice and age/sex matched WT mice were used in the current studies. Mice were maintained in a specific pathogen-free (SPF) environment with ad libitum access to food and water.

### Muscle injury

Mice were anesthetized using 2% isoflurane and muscle injury was induced by bilateral intramuscular injections of 50 μL of 1.2% barium chloride (BaCl_2_) prepared in sterile saline into the tibialis anterior (TA) muscle. Following the procedure, the mice were placed back in their respective cages and closely monitored until ambulatory. To assess the extent of inflammation and regeneration, TA muscles were collected on day 3, day 5, and day 14 following muscle injury.

### Muscle tissue collection

TA muscles were rapidly dissected under isoflurane anesthesia and muscle weights were recorded. The left TA was snap-frozen in liquid nitrogen for lipidomic analysis. The proximal portion of the right TA was snap frozen in liquid nitrogen for molecular analysis. The distal portion of the right TA was orientated longitudinal on plastic support, covered with a layer of optimal cutting temperature (OCT) compound, and then flash-frozen in isopentane chilled with liquid nitrogen. Samples were stored at −80°C until further analysis.

### Histology and immunofluorescence staining

Tissue cross-sections (10 µm) were cut from the mid-belly region of OCT embedded TA muscle using a Leica CM1950 cryostat at –20°C. Muscle tissue sections were collected onto SuperFrost Plus slides and air-dried at room temperature. Unfixed sections were used for hematoxylin and eosin (H&E) and muscle fiber type staining. Slides were fixed in acetone for 10 minutes at - 20°C before air-drying in preparation for immune cell staining. Prepared slides were blocked using either 10% normal goat serum (GS) (Invitrogen, 10000C) in phosphate buffered saline (PBS) or with Mouse on Mouse (M.O.M.) IgG Blocking Reagent (Vector Laboratories, MKB22131) when mouse primary antibodies were used on mouse tissue samples. The slides were then incubated with primary antibodies prepared in either 10% GS in PBS or M.O.M protein diluent at 4°C overnight. On the following day, the sections were washed in PBS and then incubated with Alexa Fluor-conjugated secondary antibodies (diluted 1:500 in PBS) for 1 h at room temperature. Slides were washed in PBS and then mounted with coverslips using MOWIOL Fluorescence Mounting Medium. Stitched panoramic brightfield and fluorescent images of the entire TA muscle cross section were captured using an automated fluorescent microscope (Echo Revolution) operating in upright configuration. Myofiber morphology, central nuclei fiber identification, and muscle fiber type profile was analyzed by high-throughput full automated image analysis using the MuscleJ 1.0.2 plugin for FIJI/ImageJ^31^.

### Immunohistochemistry antibodies

Primary antibodies used include MyHC type I [Developmental Studies Hybridoma Bank (DSHB), BA-D5c, 1:100], MyHC type IIA (DSHB, SC-71c, 1:100), MyHC type IIB (DSHB, BF-F3c, 1:100), eMHC (DSHB, F1.652s, 1:20), Ly6G (GR1) (Bio-Rad, MCA2387, 1:50), CD68 (Bio-Rad, MCA1957, 1:200), CD163 (Santa Cruz, sc-58965, 1:200), and laminin (Abcam, ab7463, 1:200). Primary antibody staining was visualized with appropriate Alexa Fluor conjugated secondary antibodies (Invitrogen, 1:500 in PBS). In specific experiments fluorescent dyes including DAPI (Invitrogen, Thermo Fisher Scientific, D21490, 2 μg/mL), wheat germ agglutinin (WGA) Alexa Fluor 350 conjugate (Invitrogen, W11263, 100 µg/mL), and phalloidin (Invitrogen, Thermo Fisher Scientific ActinRed 555 ReadyProbes, R37112) were used to counterstain cell nuclei, extracellular matrix, and muscle fibers, respectively.

### C2C12 Cell culture

Murine C2C12 myoblasts were sourced from the American Type Culture Collection (ATCC, CRL-1772). Myoblasts were growth at 37°C and 5% CO_2_ in high-glucose Dulbecco’s Modified Eagle’s Medium (DMEM) (Gibco, 11995073), supplemented with 10% fetal bovine serum (FBS) (Corning, 35015CV), and antibiotics [penicillin (100 U/ml) and streptomycin (100 µg/mL)] (Gibco, 15140122). Confluent myoblasts were switched to DMEM supplemented with antibiotics and 2% horse serum (HS) (Gibco, 26050088) to induce myogenic differentiation. To assess their impact on C2C12 myoblast differentiation commercially available LOX inhibitors were prepared in differentiation media and added to confluent myotube cultures at the onset of myogenic differentiation. Drugs tested included: (1) The pan LOX inhibitors 5,8,11,14-Eicosatetraynoic Acid (ETYA) (Cayman Chemicals, 90120), nordihydroguaiaretic acid (NDGA) (Cayman Chemicals, 70300), and baicalein (Cayman Chemicals, 70610). (2) The 15-LOX specific inhibitors BLX3887 (Cayman Chemicals, 27391), 9c(i472) (Cayman Chemicals, 28225), PD 146176 (Cayman Chemicals, 10010518), ML-351 (Cayman Chemicals, 16119), ThioLox (Cayman Chemicals, 38946). (3) The 5-LOX specific inhibitors malotilate (Cayman Chemicals, 30266), zileuton (Cayman Chemicals, 10006967), and MK-886 (Cayman Chemicals, 10133). (4) The 12-LOX specific inhibitors CAY10698 (Cayman Chemicals, 18582) and ML-355 (Cayman Chemicals, 18537). Following 72 h of myogenic differentiation myotube cultures were fixed in 4% paraformaldehyde (PFA) in preparation for immunocytochemistry analysis.

### Primary myoblast culture

Primary myoblasts were isolated from pooled hindlimb muscles of 6-8 month old male *Alox15*^-/-^ mice utilizing the protocol described by Hindi et al., 2017^32^. Primary myoblasts were cultivated at 37°C and 5% CO_2_ on T75 flasks coated with 10% Matrigel Matrix (Corning, 354234) in growth media (GM) consisting of Ham’s F-10 Nutrient Mixture (Gibco, 11550043), supplemented with 20% FBS, fibroblast growth factor basic (bFGF, 10 ng/mL) (PeproTech, 100-18B), and antibiotics [penicillin (100 U/ml) and streptomycin (100 µg/mL)]. To assess proliferation rate 2 × 10^5^ myoblast were plated per well of Matrigel coated 12-well plates (Thermo Scientific, 130185) and allowed to proliferate in GM for 48 h. To assess myogenic differentiation, 2 × 10^5^ myoblasts were plated per well of Matrigel coated 12-well plates and allowed to proliferate in GM for 72 h before switching to differentiation media (DM) consisting of DMEM supplemented with antibiotics and 2% horse serum (HS) (Gibco, 26050088) for a further duration of 72 h. To test the effect of LC-PUFA supplementation on myotube formation, differentiation media was supplemented with a 25 µM dose of ARA (Cayman Chemicals, 90010), EPA (Cayman Chemicals, 90110), DPA (Cayman Chemicals, 90165), DHA (Cayman Chemicals, 90310), or equimolar mixture of these four fatty acids (6.25 µM each). Following 72 h of myogenic differentiation conditioned cell culture media was collected for analysis of extracellular lipid mediator concentration by LC-MS/MS based metabolipidomic profiling and myotubes were fixed in 4% PFA for immunocytochemistry analysis.

### Immunocytochemistry and image analysis

To assess cellular morphology, myotubes were fixed in 4% PFA, permeabilized with 0.1% Triton X-100, and blocked in 1% bovine serum albumin (BSA) for 1 h at room temperature. Cells were then incubated overnight at 4°C with blocking buffer containing primary antibodies against sarcomeric MyHC (DSHB, MF20c, 1:100) and myogenin (DSHB, F5Dc, 1:100). The following morning, cells were incubated with secondary antibodies including Goat Anti-Mouse IgG2b Alexa Fluor 647 conjugate (Invitrogen, A-21242, 1:500) and Goat Anti-mouse IgG1 Alexa Fluor 555 conjugate (Invitrogen, A-21127, 1:500) for 1 h at room temperature. DAPI (Invitrogen) D21490, 2 µg/mL was used to counterstain the cell nuclei. Stained cells were visualized with an automated fluorescent microscope (Echo Revolution) operating in inverted configuration. Fluorescent images were automatically captured from a total of 9 predetermined fields of view (FOVs) per well of 12-well culture plates using a 10 × Plan Fluorite objective. To assess average myotube diameter, 50 myotubes per well were analyzed as previously described by us^33^. From each well, 5 FOVs were randomly selected, and the diameters of the 10 largest sarcomeric myosin-positive multinucleated cells in each field were manually measured at their widest uniform point using ImageJ software. For branching myocytes, each branch was measured as a separate myotube, and the region where the branches converge was exclude. To assess total myotube area per FOV, a custom in-house ImageJ macro was employed. Myosin^+^ cell area was automatically quantified for nine FOVs per culture well from 3-4 independent culture wells per experimental group.

### RNA extraction, cDNA synthesis, and RT q-PCR

Frozen TA muscle samples (10-20 mg) were homogenized for 45 seconds (4 m/s) in 600 µL of ice-cold Trizol reagent (Invitrogen, 15596018) using a Fisherbrand Bead Mill 4 mini homogenizer (Thermo Fisher Scientific, 15-340-164). RNA was extracted from muscle tissue homogenates or cellular lysates by phenol-chloroform phase separation and isopropanol precipitation following the Trizol protocol with minor modifications. The concentration and quality of RNA were assessed by spectrophotometry using a NanoDrop spectrophotometer. To eliminate any contaminating genomic DNA, the extracted RNA was treated with DNase I (Thermo Fisher Scientific, AM2222) which was then deactivated via heat treatment. RNA (1 μg) was reverse transcribed to cDNA using the High-Capacity RNA-to-cDNA Kit (Applied Biosystems, 4387406). Quantitative real-time PCR (RT-qPCR) was carried out on the cDNA samples using a QuantStudio 5 Real-Time PCR System (384-well) (Applied Biosystems, A28570). cDNA samples (1-40 ng dependent on gene of interest) were analyzed in duplicate 10 μL reaction volumes of PowerUp™ SYBR™ Green Master Mix (Applied Biosystems, A25742) with 1 μM forward and reverse primers. The relative expression levels of mRNA were evaluated using the 2^−ΔΔCT^ method. Skeletal muscle tissue, bone marrow derived MФ, and primary myoblast expression of genes of interest were normalized to *Gapdh*, *Tbp*, and *Rplp0* as internal controls, respectively. Primer sequences are listed in Table 1.

**Table 1.**
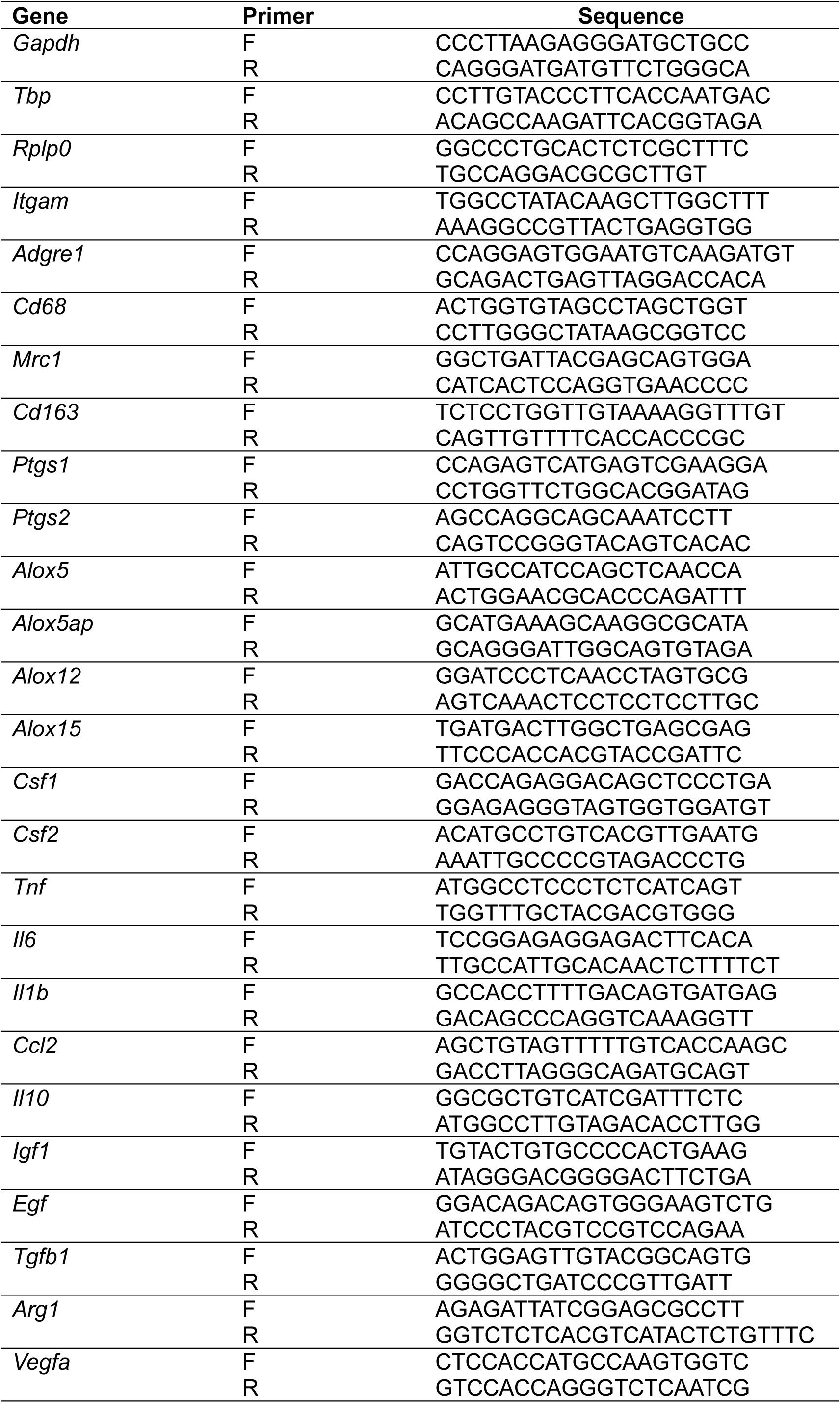
Primer sequences used for RT-qPCR.

### Protein extraction and western blotting

Frozen TA muscle samples (10-20 mg) were homogenized for 60 seconds (5 m/s) using a Fisherbrand Bead Mill 4 mini homogenizer (Thermo Fisher Scientific, 15-340-164) in ice-cold 1 × RIPA lysis buffer (MilliporeSigma, 20188) (15 µL/mg tissue) supplemented with a 1 × concentration of Halt Protease and phosphatase inhibitor cocktail (Thermo Fisher Scientific, 78442). The resulting homogenate was agitated at 4°C for 1 h and then clarified by centrifugation at 13,000 g for 10 minutes at 4°C. The supernatant was collected, and protein concentration was determined using a Pierce bicinchoninic acid (BCA) protein assay kit (Thermo Fisher Scientific, PI23225). Protein samples were diluted to a standard concentration in 1 × Laemmli buffer and boiled for 5 min. An equal volume of protein (20 µg) was then separated by sodium dodecyl sulfate-polyacrylamide gel electrophoresis (SDS-PAGE). Proteins were transferred to a nitrocellulose membrane using a Trans-Blot Turbo Transfer System (Bio-Rad, 1704150). Membranes were blocked for 1 h a room temperature in 5% skim milk power in Tris buffered saline with 0.1% Tween 20 (TBST). Membranes were subsequently incubated overnight at 4°C with gentle agitation with a Rabbit Anti-15-LOX-1 primary antibody (Abcam, ab244205) diluted 1:1000 in 5% Bovine Serum Albumin (BSA) in TBST. The following day membranes were washed in TBST and probed with a Goat Anti-Rabbit IgG (H + L) Horseradish Peroxidase (HRP) conjugated secondary antibodies (Jackson Labs, 111-035-144) diluted in 5% skim milk in TBST for 1 h at room temperature. A primary antibody against GAPDH (Santa Cruz, 32233) was used as a loading control which was detected with a Goat Anti-Mouse IgG (H + L) Horseradish Peroxidase (HRP) conjugated secondary antibodies (Jackson Labs, 115-035-146). Protein bands were visualized using Clarity Western enhanced chemiluminescent (ECL) substrate (Bio-Rad, 1705060). Chemiluminescent signals were captured using a ChemiDoc Imaging System (Bio-Rad, 12003153). Densitometry analysis was performed using Image Lab 6.1 software (Bio-Rad).

### ELISA

Conditioned cell culture media samples (cell culture supernatants) were collected from proliferating myoblasts or differentiating myotubes obtained from WT and *Alox15*^-/-^ mice. Conditioned media samples were centrifuged at 500 × g at 4°C, the supernatant collected, and stored at −80°C. Cell culture media samples were thawed once and analyzed by commercially available ELISA kit for determination of 15(S)-HETE concentration as per the manufactures’ recommendation (Cayman Chemical, 534721).

### Bone marrow-derived MФ culture

Bone marrow cells were collected from the tibias and femurs of WT and *Alox15*^-/-^ mice and cultured for 7 days at 37°C and 5% CO_2_ in MФ growth media consisting of high glucose DMEM supplemented with 10% FBS, antibiotics [penicillin (100 U/ml) and streptomycin (100 µg/mL)], and 20 ng/mL macrophage colony stimulating factor (M-CSF) (BioLegend, 576404). The resulting adherent bone marrow-derived MФ (BMMs) were then plated in MФ growth media supplemented with M-CSF at a density of 2.5 × 10^5^ cells per well of a 12-well plate (for immunocytochemistry) or 5 × 10^5^ cells per well of a 12-well plate (for RNA extraction). MФ were allowed to adhere to the plastic surface for 24 h. M-CSF was then withdrawn and BMMs were either maintained as naïve M0 MФ in serum free DMEM, polarized to M1 MФ by stimulation with 100 ng/mL LPS (Sigma-Aldrich, L2630) and 20 ng/mL interferon gamma (INF-γ) (R&D Systems, 485-MI), or polarized to M2 MФ by stimulation with 20 ng/mL interleukin-4 (IL-4) (R&D Systems, 404-ML). The resulting M0, M1, and M2 MФ were collected for RNA extraction by lysis in Trizol reagent following 24 h of polarization or fixed in 4% PFA for immunocytochemistry analysis following 48 h of polarization. Conditioned cell culture media was also collected following 24 h polarization for analysis of extracellular lipid mediators by LC-MS/MS based metabolipidomic profiling.

### LC-MS/MS–based metabolipidomic profiling of muscle tissue

TA muscle samples (20-40 mg) were mechanically homogenized in 1 mL PBS using a bead mill. The tissue homogenates were centrifuged at 3000 × g for 5 minutes and the supernatant was collected. Sample supernatants (0.85 mL) were spiked with 5 ng each of 15(S)-HETE-d8, 14(15)-EpETrE-d8, Resolvin D2-d5, Leukotriene B4-d4, and Prostaglandin E1-d4 as internal standards (in 150 μL methanol) for recovery and quantitation and mixed thoroughly. The samples were then extracted for polyunsaturated fatty acid metabolites using C18 extraction columns as previously described^21,23–25,27,34^. Briefly, the internal standard spiked samples were applied to conditioned C18 cartridges, washed with 15% methanol in water followed by hexane, and then dried under vacuum. The cartridges were eluted with 2 volumes of 0.5 mL methanol with 0.1% formic acid. The eluate was dried under a gentle stream of nitrogen. The residue was redissolved in 50 μL methanol–25 mM aqueous ammonium acetate (1:1) and subjected to LC-MS/MS analysis.

HPLC was performed on a Prominence XR system (Shimadzu) using Luna C18 (3 μm, 2.1 × 150 mm) column. The mobile phase consisted of a gradient between A, methanol-water-acetonitrile (10:85:5 v/v), and B, methanol-water-acetonitrile (90:5:5 v/v), both containing 0.1% ammonium acetate. The gradient program with respect to the composition of B was as follows: 0–1 minute, 50%; 1–8 minutes, 50%–80%; 8–15 minutes, 80%– 95%; and 15–17 minutes, 95%. The flow rate was 0.2 mL/min. The HPLC eluate was directly introduced to the electrospray ionization source of a QTRAP 5500 mass analyzer (Sciex) in the negative ion mode with following conditions: curtain gas: 35 psi, GS1: 35 psi, GS2: 65 psi, temperature: 600°C, ion spray voltage: –1500 V, collision gas: low, declustering potential: –60 V, and entrance potential: –7 V. The eluate was monitored by Multiple Reaction Monitoring (MRM) method to detect unique molecular ion–daughter ion combinations for each of the lipid mediators using a scheduled MRM around the expected retention time for each compound. Optimized collisional energies (18–35 eV) and collision cell exit potentials (7–10 V) were used for each MRM transition. Spectra of each peak detected in the scheduled MRM were recorded using enhanced product ion scan to confirm the structural identity. The data were collected using Analyst 1.7 software, and the MRM transition chromatograms were quantitated by MultiQuant software (both from Sciex). The internal standard signals in each chromatogram were used for normalization, recovery, as well as relative quantitation of each analyte.

LC-MS/MS data were analyzed using MetaboAnalyst 6.0^35^. Missing values were replaced with the estimated limit of detection (LoDs) (1/5 of the minimum positive value of each variable). Heatmaps were generated using the Euclidean distance measure and the Ward clustering algorithm following autoscaling of features without data transformation. Targeted parametric statistical analysis was also performed on a predetermined subset of metabolites of interest.

#### Muscle Force Testing

These procedures are modified from Dellorusso et al. 2001. Mice were anesthetized with 2% Isoflurane to maintain a deep anesthesia throughout the experiment. Hindlimb fur was removed with clippers. The TA muscle was exposed by removing the overlying skin and outer fasciae. The distal TA tendon was isolated and the distal half of the TA was freed from adjacent muscles by carefully cutting fasciae without damaging muscle fibers. A 4–0 silk suture was tied around the distal TA tendon, and the tendon was severed from its bony insertion. The animal was then placed on a temperature-controlled platform warmed to maintain body temperature at 37°C. A 25-gauge needle was driven through the knee and immobilized to prevent the knee from moving. The tendon was tied securely to the lever arm of a servomotor via the suture ends (6650LR, Cambridge Technology). A continual drip of saline warmed to 37°C was administered to the TA muscle to maintain its temperature. The TA muscle was stimulated with 0.2 ms pulses via the peroneal nerve using platinum electrodes. Stimulation voltage and muscle length were adjusted for maximum isometric twitch force (P_t_). While held at optimal muscle length (L_o_), the muscle was stimulated at increasing frequencies until maximum isometric tetanic force (P_o_) was reached, typically at 200 Hz, with a one-minute rest period between each tetanic contraction. Muscle length was measured with calipers, based on well-defined anatomical landmarks near the knee and the ankle. Optimum fiber length (L_f_) was determined by multiplying Lo by the TA L_f_/L_o_ ratio of 0.6^36^. After the evaluation of isometric force, the TA muscle was removed from the mouse. The tendon and suture were trimmed from the muscle, and the muscle was weighed. After removal of TA muscles, deeply anesthetized mice were euthanized by the induction of a pneumothorax. Total muscle fiber cross-sectional area (CSA) of TA muscles was calculated by dividing muscle mass by the product of Lf and 1.06 mg/mm^3^, the density of mammalian skeletal muscle^37^. Specific Po was calculated by dividing P_o_ by muscle CSA.

### Statistics

Data are presented as individual dot plots and the group means ± SEM. GraphPad Prism 10 was used for statistical analysis. Comparisons between two independent groups were performed using two-tailed unpaired t-tests, by one-way ANOVA followed by pairwise Holm-Šidák post hoc tests for three or more groups, and by two-way ANOVA followed by pairwise Holm-Šidák post hoc tests for experiments with ≥2 factors and ≥2 levels. In time-course experiments, multiple-comparisons testing was conducted using a single baseline control group. P≤0.05 were considered statistically significant.

## Results

### Leukocyte-type 12/15-LOX is expressed in skeletal muscle tissue and *Alox15*^-/-^ mice are deficient in basal intramuscular lipid mediators

Western blot analysis for the murine 12/15-LOX (15-LOX-1) protein detected a 76 kDa band in TA muscle homogenates from WT mice that matched that seen in WT spleen homogenates as a positive control (**Fig. 1A**). In contrast, *Alox15*^-/-^ mice had markedly reduced TA muscle 12/15-LOX protein expression (**Fig. 1B**). RT-qPCR also detected expression of *Alox15* mRNA in the TA muscle of WT mice, while its expression was undetectable in *Alox15*^-/-^ mice (**Fig. 1C**). LC-MS/MS analysis detected a total of 65 and 56 lipid mediators in TA muscle homogenates of WT and *Alox15*^-/-^ mice, respectively (**Table S1**). A heat map of the top 30 lipid mediators differing most between WT and *Alox15*^-/-^ mice is shown in **Fig. 1D**. Overall, 16 metabolites were significantly reduced (unadjusted p<0.05) in *Alox15*^-/-^ mice (**Table S2**). *Alox15*^-/-^ mice showed reduced intramuscular concentrations of 12/15-LOX metabolites of ARA (e.g., 12-HETE & 15-HETE), EPA (e.g., 12-HEPE), and DHA (e.g., 14-HDoHE) (**Fig. 1E-G**, **Table S2**). 15-HETE and 14-HDoHE are well-established intermediates in the biosynthesis of the lipoxin and maresin family of SPMs, respectively. However, downstream lipoxins (LXA_4_ and LXB_4_) and maresins [MaR1 and 7(S)-MaR1] were below the limits of detection of our LC-MS/MS assay. We did detect some other SPMs including RvD1, RvD6. AT-RvD3, and MaR1_n-3DPA_ **(Table S1**). But, of these only AT-RvD3 was significantly lacking in *Alox15*^-/-^ mice (**Table S2**). Interestingly, some prostaglandins with purported anti-inflammatory actions (e.g.,13,14dh-15k-PGD_2_ and PGJ_2_)^38^ were also reduced in concentrations in muscle of *Alox15*^-/-^ mice (**Table S2**). Overall, these data show that local 12/15-LOX activity is essential in maintaining a normal resting skeletal muscle lipid mediator profile.

**Figure 1:**
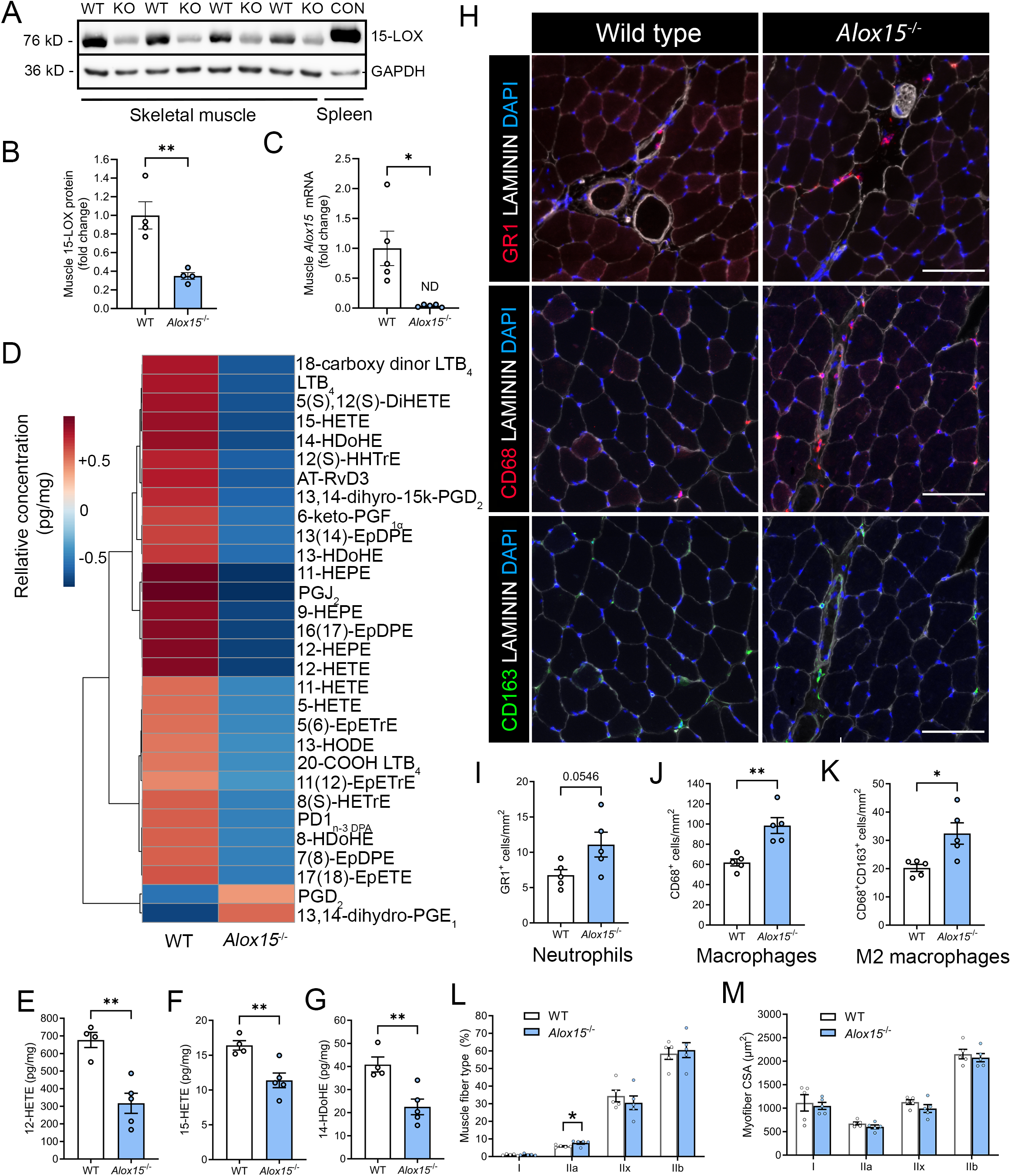
*Alox15*^-/-^ mice are deficient in intramuscular 12/15-LOX-derived lipid mediators and display chronic low-grade muscle inflammation. **A:** Western blot analysis of the abundance of the 12/15-LOX protein in TA muscle homogenates from WT and *Alox15*^-/-^ mice. A spleen homogenate from a WT mouse served as a positive control. **B:** Densitometry quantification of the abundance of the 12/15-LOX protein in the TA muscle. Protein expression was normalized to GAPDH abundance. C: *Alox15* mRNA expression in the TA muscle as determined by real-time quantitative reverse transcription PCR (RT-qPCR). Gene expression was normalized to *Gapdh*. **D:** Heatmap of the top 30 most differentially abundant lipid mediators TA muscle homogenates between WT and *Alox15*^-/-^ mice as identified by liquid chromatography-tandem mass spectrometry (LC-MS/MS). **E-G:** Quantification of the intramuscular concentration (pg/mg) of major 12/15-LOX metabolites including 12-hydroxy-eicosatetraenoic acid (12-HETE) (**E**), 15-hydroxy-eicosatetraenoic acid (15-HETE) (**F**), and 14-hydroxy-docosahexaenoic acid (14-HDoHE) (**G**) in TA muscle homogenates as determined by LC-MS/MS. **H:** TA muscle cross-sections were stained with primary antibodies against the neutrophil marker GR1, the pan monocyte/macrophage marker CD68, or the M2 macrophage marker CD163. Scale bars are 100 µm. **I-K:** Quantitative analysis of intramuscular numbers of neutrophils (GR1^+^ cells) (**I**), monocyte/macrophages (CD68^+^ cells) (**J**), and M2-like macrophages (CD68^+^CD163^+^ cells) (**K**). **L-M:** Quantification of percentage TA muscle fiber type (**L**) and fiber type specific myofiber cross-sectional area (CSA) (**M**) as determined by MuscleJ 1.0.2 software. Bars show the mean ± SEM of 4-5 mice per group (biological replicates) with dots representing data from each individual mouse. P*-*values were determined by two-tailed unpaired t-tests. *p<0.05, **p<0.01 for WT vs. *Alox15*^-/-^ mice.

### Basal skeletal muscle phenotype in WT and *Alox15*^-/-^ mice

To examine the potential consequences of a lack of 12/15-LOX derived lipid mediators, muscle mass and strength were compared between WT and *Alox15*^-/-^ mice. *Alox15* knockout had no effect on body weight (**Fig. S1A**), absolute *in situ* TA muscle strength (**Fig. S1B**), or muscle strength when normalized to muscle size (maximal specific force) (**Fig. S1C**). Finally, although the mass of the quadriceps (QUAD) muscle was significantly reduced in *Alox15*^-/-^ mice, there was no such difference in mass of the other hind-limb muscles including the soleus (SOL), plantaris (PLA), tibialis anterior (TA), or gastrocnemius (GAST) (**Fig. S1D**). Overall, these data suggests that the development and maintenance of skeletal muscle tissue appears generally undisturbed in young adult whole-body 12/15-LOX knockout mice.

### *Alox15*^-/-^ mice show chronic low-grade skeletal muscle inflammation

Some 12/15-LOX pathway metabolites (e.g., SPMs) have been implicated as playing an important counterregulatory role in the inflammatory response^20^. Therefore, we sought to quantify leukocyte numbers in TA muscles obtained from WT and *Alox15*^-/-^ mice (**Fig. 1H**). *Alox15*^-/-^ mice displayed a statistical trend (p=0.055) toward greater numbers of intramuscular PMNs (GR1^+^ cells) (**Fig. 1I**), as well as more total MФ (CD68^+^ cells) (**Fig. 1J**), and M2-like MФ (CD68^+^CD163^+^ cells) (**Fig. 1K**). Analysis of TA muscle fiber type profile revealed a higher proportion of type IIA myofibers in *Alox15*^-/-^ mice, but there was no difference in the percentage of type I, IIX, or IIB fibers. (**Fig 1L**). *Alox15*^-/-^ mice also did not differ significantly from WT mice in the mean cross-sectional area (CSA) of type I, IIA, IIX, or IIB myofibers (**Fig. 1M**). Representative fiber type staining is shown in **Fig. S1E**. Overall, these data show that a deficiency in *Alox15* results in chronic low-grade muscle inflammation and a shift in fiber type profile, but no overt evidence of myofiber atrophy.

### *Alox15*^-/-^ mice mount an exaggerated innate immune response to acute skeletal muscle injury

To assess the impact of 12/15-LOX deficiency on the acute inflammatory response to myofiber injury, TA muscles were collected at day 3 (D3), day 5 (D5), and day 14 (D14) following intramuscular injection of barium chloride (BaCl_2_). Muscle mRNA expression of the pan myeloid cell marker CD11b (*Itgam*) (**Fig 2A**), the MФ marker F4/80 (*Adgre1*) (**Fig 2B**), and the M2-like MФ marker CD206 (*Mrc1*) (**Fig. 2C**) increased markedly (>100-fold) on D3 and were each relatively greater in *Alox15^-/-^*vs. WT mice (**Fig 2A-C).** Local expressions of hematopoietic growth factors including macrophage colony-stimulating factors (M-CSF, *Csf1*) (**Fig. 2D**) and granulocyte-macrophage colony-stimulating factor (GM-CSF, *Csf2*) (**Fig. 2E**) also increased greatly on D3. Furthermore, GM-CSF, but not M-CSF, was also substantially greater in *Alox15*^-/-^ vs WT mice at this time-point (**Fig 2E**). 12/15-LOX knockout did not affect the induction of a range of other classical pro- and anti-inflammatory cytokines such as TNFα (*Tnf1*) (**Fig. S2A**), IL-1β (*Il1b*) (**Fig. S2B**), IL-6 (*Il6*) (**Fig. S2C**), or IL-10 (*Il10*) (**Fig. S2D**). Nevertheless, monocyte chemoattractant protein 1 (MCP-1, *Ccl2*) was significantly lower in *Alox15*^-/-^ vs. WT mice on D3 (**Fig. S2E**). Interestingly, expression of insulin like growth factors (IGF-1, *Igf1*) (**Fig. S2F**), transforming growth factor beta (TGF-β, *Tgfb1*) (**Fig. S2H)**, and arginase-1 (ARG-1, *Arg1*) (**Fig. S2I)** were each significantly greater in *Alox15*^-/-^ mice on D3. On the other hand, vascular endothelial growth factor (VEGF, *Vegfa*) was blunted in *Alox15*^-/-^ mice on D14 relative to WT controls (**Fig. S2J**).

**Figure 2:**
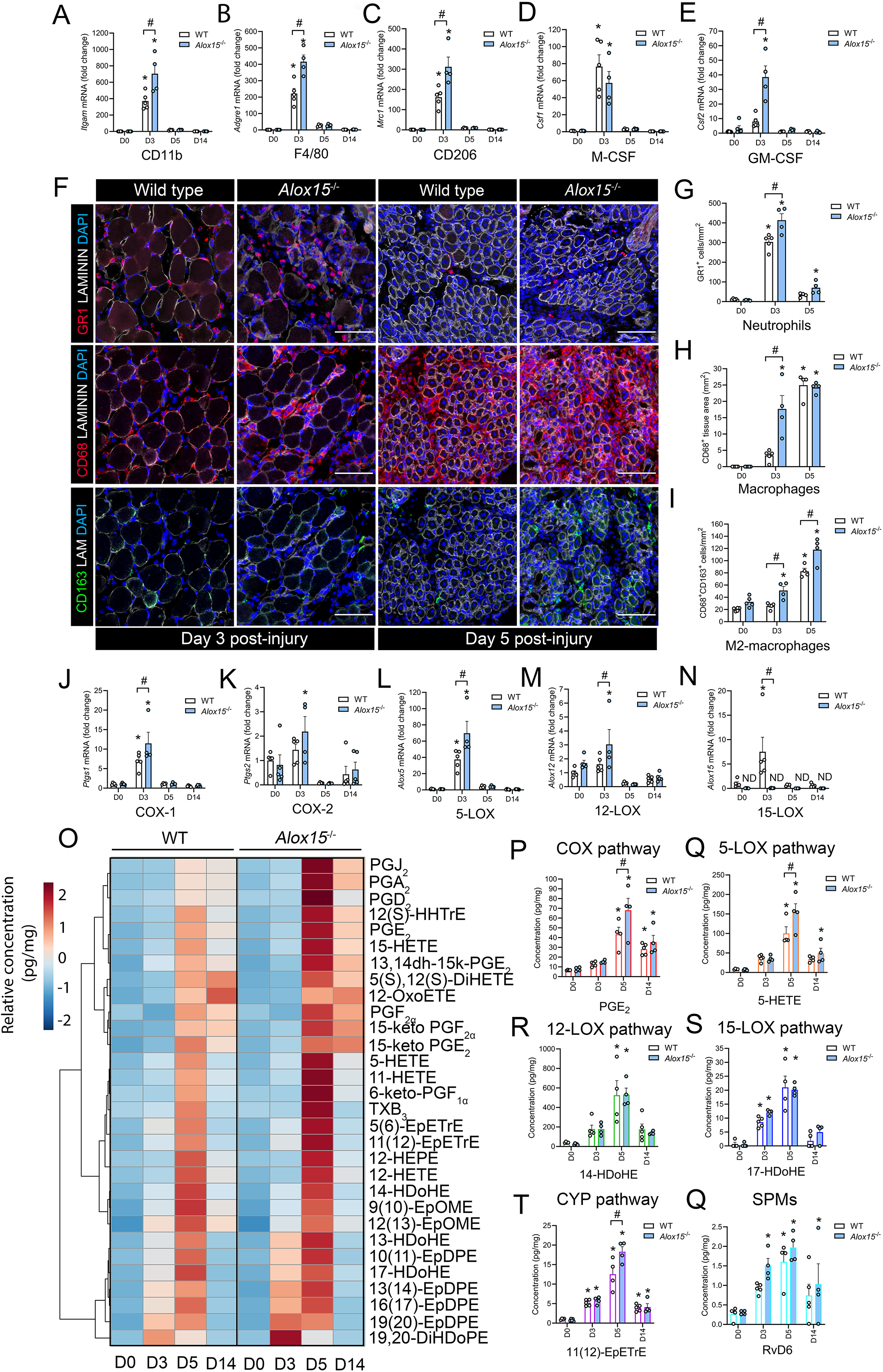
Leukocyte-type 12/15-LOX deficient mice display greater local inflammation and an imbalance of pro-inflammation vs. anti-inflammatory/pro-resolving lipid mediators following acute skeletal muscle injury. **A-E:** Tibialis anterior (TA) muscle mRNA expression of the pan myeloid cell marker CD11b (*Itgam*) (**A**), the pan monocyte/ MФ marker F4/80 (*Adgre1*) (**B**), the M2 MФ marker CD206 (*Mrc1*) (**C**), the hematopoietic growth factor M-CSF (*Csf1*) (**D**), and the hematopoietic growth factor GM-CSF (*Csf2*) (**E**) in wild type (WT) and *Alox15*^-/-^ mice at day 0 (D0), day 3 (D3), day 5 (D5), and day 14 (D14) following myofiber injury induced by intramuscular injection of 50 µL of 1.2% barium chloride (BaCl_2_). Expression of genes of interest was normalized to *Gapdh*. **F:** Immunofluorescence staining with primary antibodies against neutrophils (PMNs) (GR1), monocytes/MФ (CD68), and M2 MФ (CD163) in injured TA muscle of WT and *Alox15^-/-^* mice on D3 and D5 post-injury. Cell nuclei were stained with DAPI and a primary antibody against laminin was used to identify muscle fiber boundaries. Scale bars are 100 µm. **G-I**: Quantification of PMNs (GR1^+^ cells/mm^2^) (**G**), monocytes/ MФ (CD68^+^ cells/mm^2^) (**H**), and M2 MФ (CD68^+^CD163^+^ cells/mm^2^) (**I**) at D0, D3, and D5 post-injury in WT and *Alox15^-/-^*mice. **J-N:** TA mRNA expression of major lipid mediator biosynthesis enzymes including COX-1 (*Ptgs1*) (**J**), COX-2 (*Ptsg2*) (**K**), 5-LOX (*Alox5*) (**L**), 12-LOX (*Alox12*) (**M**), and 15-LOX (*Alox15*). **O:** Heatmap of the top 30 most differentially abundant lipid mediators between WT and *Alox15*^-/-^ mice as identified by liquid chromatography-tandem mass spectrometry (LC-MS/MS) analysis of TA muscle homogenates. **P-Q:** Quantification of major representative metabolites of the COX pathway (e.g., PGE_2_) (**P**), 5-LOX pathway (e.g., 5-HETE) (**Q**), 12-LOX pathway (e.g., 14-HDoHE) (**R**), 15-LOX pathway (e.g., 17-HDoHE) (**S**), CYP pathway [e.g., 11(12)-EpETrE) (**T**), and downstream bioactive SPMs (e.g., RvD6) (**Q**). Bars show the mean ± SEM of 4-5 mice per group (biological replicates) with dots representing data from each individual mouse. P*-*values were determined by two-way ANOVA followed by Holm-Šídák post-hoc tests. *p<0.05 vs. D0 and #p<0.05 for WT vs. *Alox15^-/-^* mice.

Immunofluorescent staining revealed that muscle infiltration of blood PMNs (GR1^+^ cells) peaked on D3 post-injury (**Fig 2F**). Additionally, *Alox15^-/-^* mice showed relatively greater intramuscular numbers of PMNs at this time-point (**Fig 2G**). Intramuscular infiltration of MФ (CD68^+^ cells) peaked on D5 (**Fig 2F**). When compared to WT controls, *Alox15^-/-^* mice showed an exaggerated CD68^+^ cell infiltration at D3, but similar intramuscular numbers of MФs by D5 (**Fig. 2H**). M2 MФ (CD68^+^CD163^+^ cells) numbers also peaked on D5 and were more numerous in *Alox15^-/-^* mice at both D3 and D5 (**Fig. 2I**). These data show that *Alox15*^-/-^ mice exhibit an overall relatively greater acute inflammatory response when compared to WT mice.

### Dysregulated local lipid mediator response to skeletal muscle injury in *Alox15^-^*^/-^ mice

RT-qPCR analysis revealed that muscle mRNA expression of major lipid mediator biosynthetic enzymes including COX-1 (*Ptgs1*) (**Fig. 2J**), 5-LOX (*Alox5*) (**Fig. 2L**), 5-LOX activating protein (FLAP, *Alox5ap*) (**data not shown**), and 15-LOX (*Alox15*) (**Fig 2N**) all increased greatly on D3. As expected, *Alox15* mRNA expression was undetectable in *Alox15*^-/-^ mice at D3, D5, and D14 (**Fig 2N**). On the other hand, COX-1 (*Ptgs1*) (**Fig. 2J**), COX-2 (*Ptgs2*) (**Fig. 2K**), 5-LOX (*Alox5*) (**Fig. 2L**), FLAP (*Alox5ap*) (**data not shown**), and 12-LOX (*Alox12*) (**Fig. 2M**) were each more highly expressed in *Alox15*^-/-^ vs. WT mice on D3. Overall, these data show that *Alox15*^-/-^ mice appear to mount a compensatory transcriptional upregulation of other lipid mediator biosynthetic enzyme in the absence of leukocyte-type 12/15-LOX.

LC-MS/MS-based metabolomic profiling revealed that a total of 53, 72, and 48 lipid mediators increased (unadjusted p<0.05) in muscle tissue of WT mice at D3, D5, and D14 post-injury, respectively (**Table S3**). In contrast, a total of only 5, 0, and 2 analytes decreased in muscle tissue of WT mice at D3, D5, and D14, respectively (**Table S3**). A heatmap of the top 30 lipid mediators most influenced by BaCl_2_-induced muscle damage is shown in **Fig. 2O**. Following muscle injury there was increased local concentrations of many LC-PUFA metabolites of the COX pathway (e.g., PGE_2_), 5-LOX pathway (e.g., 5-HETE), 12-LOX pathway (e.g., 14-HDoHE), 15-LOX pathway (e.g., 17-HDoHE), and CYP pathway [e.g., 5(6-EpETrE)] (**Fig. 2O**). Some downstream SPMs including LXA_4_, RvD1, RvD2, and RvD6 were also detected at increased concentrations in regenerating muscle tissue on D5 and/or D14 (**Table S3**). When compared to WT controls, *Alox15*^-/-^ mice showed relatively greater concentrations of many n-6 ARA metabolites of the COX pathway (e.g., PGE_2_) (**Fig. 2P & Fig S3A**), the 5-LOX pathway (e.g., 5-HETE) (**Fig. 2Q & Fig S3B**), and the CYP pathway (e.g., [11(12)-EpETrE]) (**Fig. 2T & Fig. S3E**). In contrast, major n-3 DHA metabolites of 12-LOX pathway (e.g., 14-HDoHE) (**Fig. 2R & Fig S3C**) and 15-LOX pathway (e.g., 17-HDoHE) (**Fig. 2S & Fig. S3D**) were surprisingly detected at similar concentrations in WT and *Alox15*^-/-^ mice. Similarly, downstream n-3 series SPMs (e.g., RvD6) did not differ significantly between WT and *Alox15*^-/-^ mice (**Fig. 2Q & Fig S3F**). Surprisingly, concentrations of the n-6 ARA derived SPM LXA_4_ were significantly greater in *Alox15*^-/-^ vs. WT mice on D5 (**Fig. S2F**), as were levels of its biosynthetic intermediate 15-HETE (**Fig. S2D**). While E-series resolvins (e.g., RvE1) were not detected, the E-resolvin pathway marker 18-HEPE was also significantly greater in *Alox15*^-/-^ vs WT mice on D5 (**Fig. S3F**). Overall, these data reveal an absolute overabundance of pro-inflammatory n-6 metabolites of the COX and 5-LOX pathways, together with a relative deficiency of n-3 DHA derived anti-inflammatory/pro-resolving mediators of the 12/15-LOX pathways in *Alox15*^-/-^ mice. Nevertheless, some potential n-6 PUFA metabolites of the 15-LOX pathway, such as 15-HETE and LXA_4_ are greater in *Alox15*^-/-^ mice.

### Deleterious effects of *Alox15* deficiency on skeletal muscle regeneration

To investigate whether 12/15-LOX deficiency might impact upon the efficiency of skeletal muscle regeneration we first measured mRNA expression of key myogenic genes throughout the time-course of muscle injury and regeneration. *Alox15*^-/-^ mice showed significantly lower basal muscle expression of myogenic differentiation 1 (MyoD, *Myod1*) (**Fig. 3A**) and myogenic regulatory factor 4 (MRF4, *Myf6*) (**Fig. 3D**), but similar expression of myogenic factor 5 (MYF5, *Myf5*) (**Fig. 3B**) and myogenin (MyoG, *Myog*) (**Fig. 3C**). Expression of *Myod1*, *Myf5*, and *Myog* each increased greatly on day 5 post-injury, while *Myf6* mRNA rather decreased far below baseline levels at this time-point. When compared to WT controls, expression of *Myog* was significantly blunted on day 5 post-injury in *Alox15*^-/-^ mice (**Fig. 3C**). On day 14 post-injury, muscle expression of *Myf6* was also lower in *Alox15*^-/-^ mice (**Fig. 3D**) and a similar statistical trend was also observed for *Myod1* (**Fig. 3A**). Muscle mRNA expression of the developmental embryonic myosin isoform (eMHC, *Myh3*) increased markedly on D5, and this response was also significantly diminished in *Alox15*^-/-^ mice (**Fig. 3E**). Overall, these data suggest overall suppressed local induction of myogenic genes in *Alox15*^-/-^ mice during the regenerative phase following acute muscle injury.

**Figure 3:**
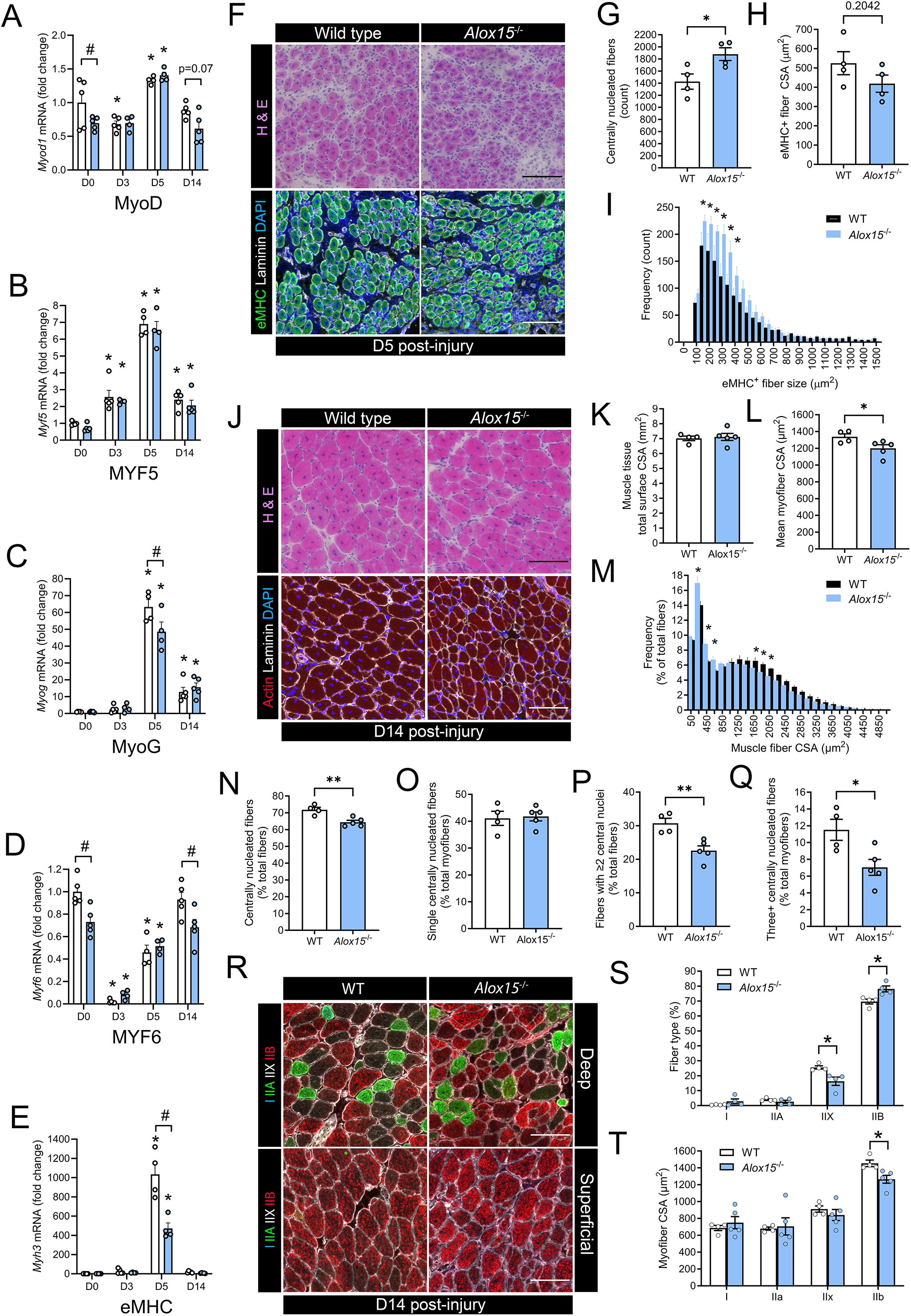
Deleterious effect of leukocyte-type 12/15-LOX deficiency on skeletal muscle regeneration: **A-E** Muscle mRNA expression of myogenic genes including MyoD (*Myod1*) (**A**), MYF5 (*Myf5*) (**B**), MyoG (*Myog*) (**C**), MYF6 (*Myf6*) (**D**), and eMHC (*Myh3*) **(E)** at day 3 (D3), day 5 (D5), and day 14 (D14) following muscle injury induced by intramuscular injection of BaCl_2_. Expression of genes of interest was normalized to *Rplp0.* **F:** Cross-sections of tibialis anterior (TA) muscles from wild type (WT) and *Alox15*^-/-^ mice obtained on D5 post-injury were stained with Hematoxylin & Eosin (H & E) or with primary antibodies against eMHC, laminin, and DAPI to assess regenerating muscle fiber morphology. Scale bars are 100 µm. **G-I:** Quantitative analysis of the absolute number of regenerating (centrally nucleated) myofibers (**G**), mean regenerating (eMHC^+^) myofiber cross-sectional area (CSA) (**H**), and frequency distribution of regenerating (eMHC+) myofiber CSA (**I**). **J:** TA muscle sections obtained on D14 post-injury were stained with H & E or with primary antibodies against actin, laminin, and DAPI for analysis of regenerating muscle fiber morphology. Scale bars are 100 µm. **K-M:** Quantification of the total TA muscle cross-sectional area (CSA) (**K**), mean TA muscle fiber CSA (**L**), and frequency distribution of myofiber CSA (**M**) within TA muscle cross-sections obtained from WT and *Alox15*^-/-^ mice on D14 post-injury. **N-Q:** Quantification of the percentage of centrally nucleated myofibers (**N**), and the proportion of regenerating muscle fibers containing one (**O**), two (**P**), or ≥three (**Q**) centrally located myonuclei in TA muscle cross-sections from WT and *Alox15*^-/-^ mice on D14 post-injury. **R:** TA muscle cross-sections obtained on day 14 post-injury were stained with primary antibodies against type I, IIA, and IIB myosin heavy chain (MyHC). Type IIX myofibers remain unstained (black). Scale bars are 100 µm. **S-T:** Quantification of percentage muscle fiber type profile (**S**) and fiber type specific myofiber CSA (**T**). Bars show the mean ± SEM of 4-5 mice per group (biological replicates) with dots representing data from each individual mouse. **A-E:** P*-*values were determined by two-way ANOVA followed by Holm-Šídák post-hoc tests. *p<0.05 vs. D0 and #p<0.05 for WT vs. *Alox15^-/-^* mice. **F-T:** P*-*values were determined by two-tailed unpaired t-tests. *p<0.05, **p<0.01 for WT vs. *Alox15*^-/-^ mice.

Histological analysis revealed that both WT and *Alox15^-/-^*mice mounted a robust regenerative response on D5 as indicated by a predominance of many small myofibers with centrally located myonuclei that expressed eMHC (**Fig. 3F**). At this point, *Alox15*^-/-^ mice showed more numerous regenerating myofibers (**Fig. 3G**). Nevertheless, these fibers tended to be relatively smaller in size when compared to WT mice (**Fig. 3H**). Indeed, frequency distribution analysis showed many more eMHC^+^ myofibers with a CSA in the range of 150-400 µm^2^ in *Alox15*^-/-^ vs. WT mice (**Fig 3I**). Regenerating myofiber size increased markedly (>2-fold) in both WT and *Alox15*^-/-^ mice between D5 and D14 (**Fig. 3J**). On D14 TA muscles from *Alox15^-/-^* mice were like WT in overall size (**Fig. 3K**). Nevertheless, the average myofiber CSA was significantly lower in *Alox15^-/-^* mice (**Fig. 3L**). This was attributable to both a relatively greater proportion of very small myofibers and lower proportion of larger myofibers in *Alox15^-/-^* mice (**Fig. 3M**). The overall proportion of centrally nucleated myofibers was also significantly reduced in *Alox15^-/-^* mice at D14 (**Fig 3N**). Interestingly, there was no difference between groups in the proportion muscle fibers containing a single centrally located myonuclei (**Fig. 3O**). However, the proportion of more mature regenerating myofibers with two central nuclei (**Fig. 3P**), as well as those with ≥3 central nuclei (**Fig. 3Q**) were significantly reduced in *Alox15^-/-^* mice. Muscle fiber type staining showed that, as expected, regenerating muscles of both WT and *Alox15*^-/-^ were predominantly comprised of fast twitch type II myofibers (IIB, IIX, and IIA) (**Fig. 3R**). When compared to WT mice, regenerating muscles from *Alox15*^-/-^ mice contained a relatively increased proportion of type IIB myofibers together with a corresponding reduction in the proportion of type IIX myofibers (**Fig. 3S**). Furthermore, the mean CSA of the type IIB myofiber population, which comprised most TA muscle fibers, was significantly reduced in *Alox15*^-/-^ mice (**Fig. 3T**). Overall, these data show that *Alox15^-/-^* mice mount an impaired cellular and molecular regenerative response to muscle injury.

### *Alox15*^-/-^ MФ show heightened cytokine expression, defective M2 polarization, and an altered lipid mediator profile

We next investigated whether there may be inherent differences in the innate immune cell response that might contribute to poor muscle regeneration outcomes in *Alox15*^-/-^ mice. Naïve (M0) bone marrow-derived MФ (BMMs) were cultured *in vitro* and then polarized into pro-inflammatory M1 or anti-inflammatory M2 subsets by exposure to LPS + INF-γ or IL-4, respectively. RT-qPCR showed that expression of pro-inflammatory cytokines including IL-1β (*Il1b*) (**Fig. 4A**) and IL-6 (*Il6*) (**Fig. 4B**) were greatly increased following M1 polarization and that *Alox15*^-/-^ M1 MФ expressed even higher levels of these cytokines than WT cells (**Fig. 4A & B**). Other genes induced by M1 polarization, but not differing between WT and *Alox15^-/-^* BMMs, included F4/80 (*Adgre1*), MCP-1 (*Ccl2*), TNFα (*Tnf)*, M-CSF (*Csf1*), IL-10 (*Il10*) (**Fig S4**). Anti-inflammatory MФ markers including CD206 (*Mrc1)* (**Fig. 4C**) and annexin A1 (ANXA1) (**Fig. 4D**) were markedly upregulated following M2 polarization and these responses were blunted in *Alox15*^-/-^ cells. Expression of CD68 (*Cd68*) was also increased by M2 polarization, but did not differ between WT and *Alox15*^-/-^ BMMs (**Fig. S4**). Immunofluorescence staining also showed that *Alox15*^-/-^ MФ were less responsive to M2 polarization than WT MФ based on protein expression of anti-inflammatory MФ cell surface markers including CD163 and CD206 (**Fig. 4E & F**).

**Figure 4:**
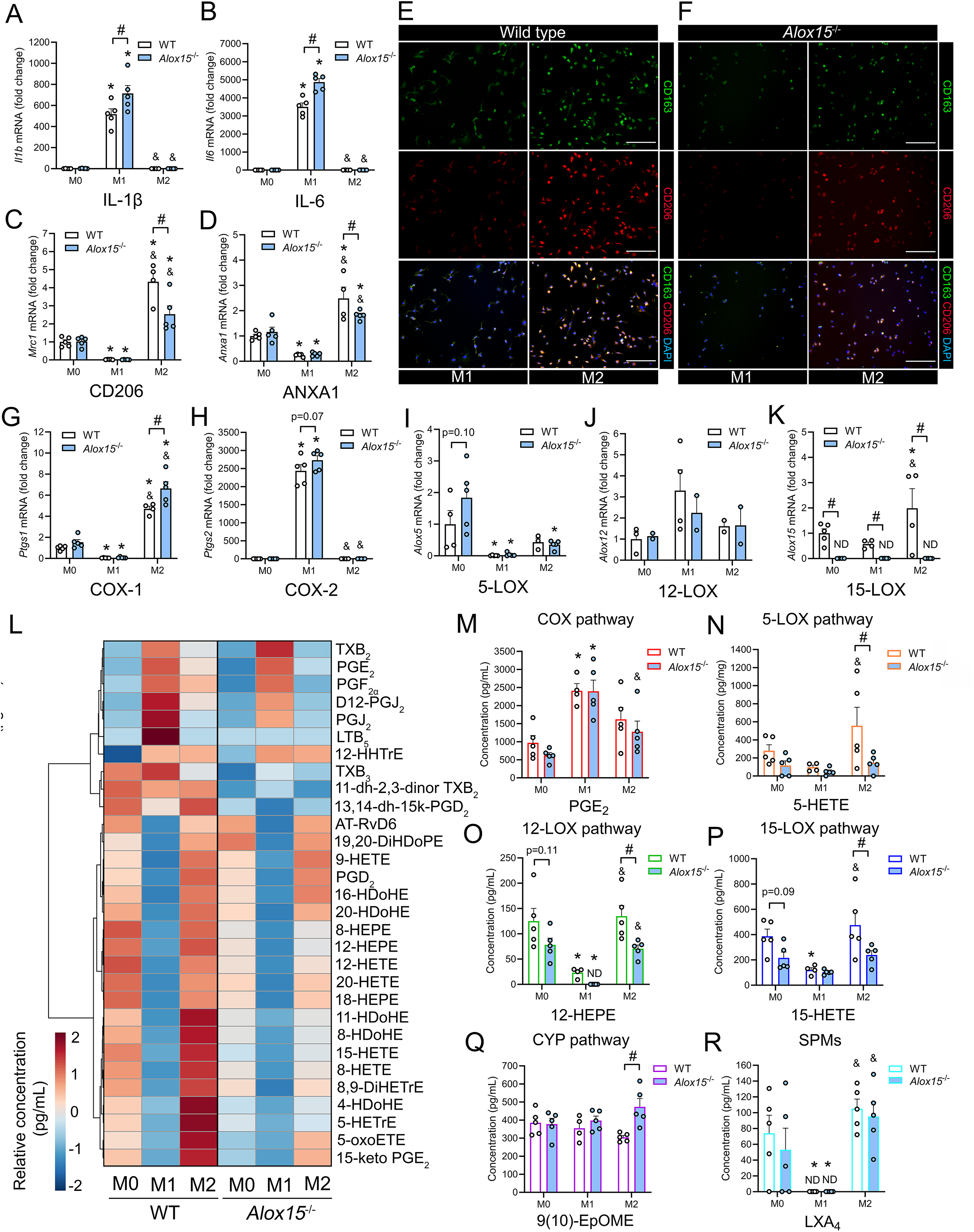
Bone marrow derived-macrophages obtained from *Alox15*^-/-^ mice display shifts in secreted lipid mediator profile and impaired M2 polarization. Bone-marrow-derived macrophages (MФ) (BMMs) obtained from wild type (WT) and 12/15-LOX deficient (*Alox15*^-/-^) mice were maintained in serum free media as naïve M0 MФ, or induced to differentiate into either M1 MФ by exposure to lipopolysaccharide (LPS) (100 ng/mL) and interferon gamma (IFN−*γγ*) (20 ng/mL), or M2 MФ by exposure to interleukin-4 (IL-4) (20 ng/mL). Following 24 h of polarization mRNA expression levels of pro-inflammatory cytokines including interleukin-1β (*Il1b*) (**A**) and interleukin-6 (*Il6*) (**B**), or M2 MФ markers including CD206 (*Mrc1*) (**C**) and annexin A1 (*Anxa1*) (**D**) were measured by real-time quantitative reverse transcription PCR (RT-qPCR). **E-F:** BMMs from WT mice (**E**) and *Alox15*^-/-^ mice (**F**) were polarized to a M1 or M2 activation state for 48 h, fixed in 4% paraformaldehyde (PFA), and stained with primary antibodies against CD163 (green) and CD206 (red). Nuclei were counterstained by DAPI (blue). Scale bars are 200 µm. **G-K:** Gene expression of major lipid mediator biosynthesis enzymes including COX-1 (*Ptgs1*) (**G**), COX-2 (*Ptsg2*) (**H**), 5-LOX (*Alox5*) (**I**), platelet-type 12-LOX (*Alox12*) (**J**), and 15-LOX-1 (*Alox15*) (**K**) in M0, M1, and M2 MФ obtained from WT and *Alox15*^-/-^ mice following 24 h of polarization. **L:** A heatmap of the top 30 most differentially modulated lipid metabolites detected in conditioned cell culture media samples obtained from WT and *Alox15*^-/-^ M0, M1, and M2 MФ following 24 h incubation in serum free DMEM as assessed by targeted liquid chromatography-tandem mass spectrometry (LC-MS/MS). **M-R:** Concentrations of major representative lipid mediator metabolites of the cyclooxygenase (COX) pathway (e.g., PGE_2_) (**M**), 5-lipoxygenase (5-LOX) pathway (e.g., 5-HETE) (**N**), 12-lipoxygenase (12-LOX) pathway (e.g., 12-HEPE) (**O**), 15-lipoxygenase (15-LOX) pathway (e.g., 15-HETE) (**P**), cytochrome P450 (CYP 450) pathway [e.g., (9(10)-EpOME)] (**Q**), and downstream bioactive SPMs (e.g., LXA_4_) (**R**). Bars show the mean ± SEM of cells obtained from 5 mice per group (biological replicates) with dots representing BMMs from each individual mouse. P*-*values were determined by two-way ANOVA followed by Holm-Šídák post-hoc tests. *p<0.05 vs. M0, &p<0.05 vs. M1, and #p<0.05 for WT vs. *Alox15^-/-^* cells.

We next assessed the impact of polarization and 12/15-LOX knockout on lipid mediator biosynthesis pathways in BMMs. M1 polarization markedly increased MФ mRNA expression of *Ptgs2* (COX-2) (**Fig. 4H**), while reducing expression of *Ptgs1* (COX-1) (**Fig. 4G**), *Alox5* (5-LOX) (**Fig. 4I**), and Alox5ap (FLAP) (**Fig. S4I**). In contrast, M2 polarization increased MФ mRNA expression of *Ptgs1* (COX-1) (**Fig. 4G**), *Alox15* (15-LOX) (**Fig. 4K**), and Alox5ap (FLAP) (**Fig. S4H**), while reducing expression of *Ptgs2* (COX-2) (**Fig. 4H**) and *Alox5* (5-LOX) (**Fig. 4I**). Expression of *Alox12* (platelet-type 12-LOX) was not affected by MФ polarization (**Fig. 4J**). As expected, *Alox15* mRNA expression was undetected in *Alox15*^-/-^ M0, M1, and M2 MФ (**Fig. 4K**). *Alox15*^-/-^ BMMs expressed significantly higher levels of *Ptgs1* (COX-1) under M2 conditions (**Fig. 4G**), tended to express higher levels of *Ptgs2* (COX-2) under M1 conditions (**Fig. 4H**), and tended to express higher *Alox5* (5-LOX) under M0 conditions (**Fig. 4I**).

LC-MS/MS profiling of serum free conditioned cell culture media samples revealed that a total of 36 lipid mediators were altered following 24 h of polarization of WT MФ from a M1 to M2 phenotype (**Table S5**). A further 18, 17, and 26 lipid mediators differed between WT and *Alox15*^-/-^ MФ under identical M0, M1, and M2 conditions (**Table S6**). The top 30 extracellular lipid mediators influenced by polarization and/or *Alox15* deficiency are shown in **Fig. 4L**. Naïve M0 MФ from *Alox15*^-/-^ mice produced relatively lower concentrations than WT of some 12/15-LOX metabolites of n-6 ARA [e.g., 15-HETE, 15-Oxo-ETE, and 5(S),12(S)-DiHETE] and n-3 DHA (e.g., AT-RvD3 and RvD5) (**Table S6**). M1 polarization greatly increased extracellular concentrations of pro-inflammatory COX metabolites of n-6 ARA (e.g., PGE_2_) (**Fig. 4M**), **Fig S4A**). An exception was PGD_2_ which was markedly reduced by M1 polarization (**Fig S4A**). Prostaglandin biosynthesis was generally similar in WT and *Alox15*^-/-^ cells under M1 conditions (**Fig. 4L**). Nevertheless, *Alox15*^-/-^ M1 MФ did produce lower amounts of PGI_2_ (measured as 6-keto-PGF_1α_) than WT cells (**Table S6**). In WT MФ, M2 polarization decreased production of most COX metabolites (**Fig S5A**), while increasing production of many potential metabolites of the 5-LOX pathway (e.g., 5-HETE) (**Fig. 4N, Fig S5B**), the 12-LOX pathway (e.g., 12-HETE) (**Fig. 4O, Fig S4C**), and the 15-LOX pathway (e.g., 15-HETE) (**Fig. 4P, Fig S5D**). Production of many of these LOX metabolites was significantly diminished in *Alox15*^-/-^ MФ (**Fig. 4N-P, Fig S5**). M2 polarization of WT MФ also increased production of several di-and tri-hydroxy LOX metabolites (e.g., the SPMs) including LXA_4_, PD1, AT-PD1, AT-RvD6, and MaR1_n-3DPA_ (**Fig 4R**, **Fig. S5F**). However, of these only AT-PD1 and MaR1_n-3DPA_ were significantly lower concentrations in conditioned media samples obtained from *Alox15*^-/-^ MФ. While most lipid mediators were diminished by 12/15-LOX deficiency, *Alox15*^-/-^ M2 MФ produced greater amounts of some CYP pathway metabolites of linoleic acid (LA) [e.g., 9(10-EpOME)] (**Fig. 4Q, Fig S5E**). Overall, these data show that M1 polarization results in a COX dominated pro-inflammatory lipid mediator profile while M2 polarization leads to a shift to a LOX dominated anti-inflammatory/pro-resolving lipid signature. Furthermore, *Alox15*^-/-^ MФ produce lower amounts of many LOX-derived monohydroxy-PUFA metabolites under M2-polarizing conditions.

### Skeletal muscle cells secrete autocrine/paracrine 12/15-LOX metabolites that directly stimulate *in vitro* skeletal muscle cell growth and development

We next examined whether muscle cells might also themselves express *Alox15* and whether a local deficiency of 12/15-LOX might directly influence muscle cell growth and development independent of immune-muscle cell crosstalk. Primary myoblasts were isolated from WT and *Alox15^-/-^* mice and cultured *in vitro*. RT-qPCR detected the expression of *Alox15* mRNA in both WT myoblast and myotube cultures (**Fig. 5A**). In contrast, *Alox15* mRNA expression was undetectable in myoblast or myotube cultures obtained from *Alox15*^-/-^ mice (**Fig. 5A**). ELISA analysis of conditioned culture media samples revealed readily detectable extracellular concentrations of the 15(S)-HETE in conditioned culture media samples obtained from WT myoblasts and myotubes, but 15(S)-HETE was significantly reduced in conditioned media samples obtained from *Alox15^-/-^* cells (**Fig. 5B**).

**Figure 5:**
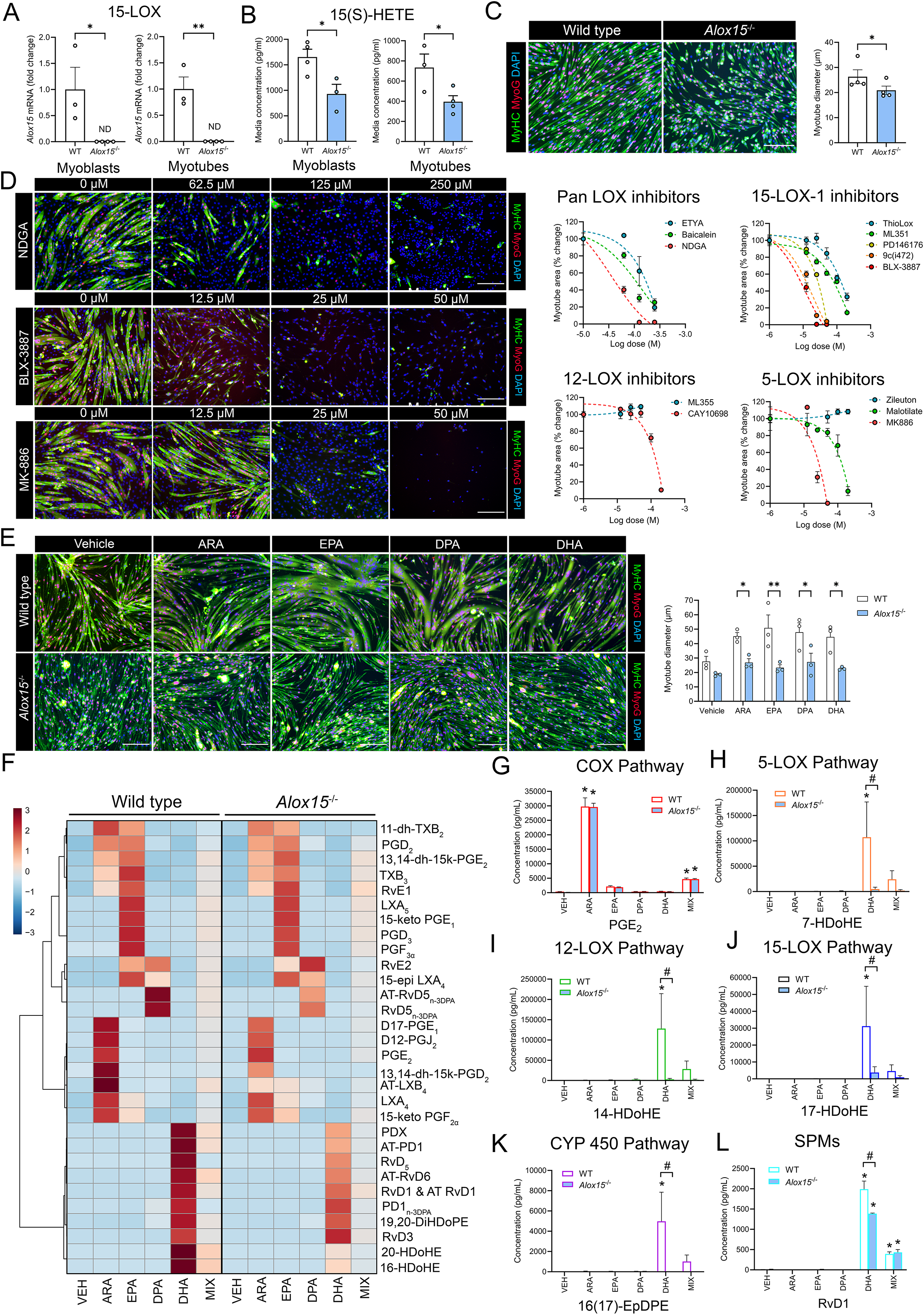
Leukocyte-type 12/15-LOX is a novel and direct determinant of myogenic progenitor cell fate. **A-B:** Primary myoblasts derived from wild type (WT) and *Alox15^-/-^*mice were cultured *in vitro* and then induced to differentiate into myotubes. The expression level of *Alox15* mRNA was measured by RT-qPCR (**A**). The production of 15(S)-HETE in conditioned culture media obtained from primary myoblasts and myotubes was also quantified by ELISA (**B**). **C:** WT and *Alox15*^-/-^ myotubes were stained for myosin heavy chain (green) and myogenin (red) following 72 h of myogenic differentiation. Nuclei were counterstained by DAPI (blue). Mean myotube diameter was measured to assess the effect of *Alox15* deficiency on skeletal muscle cell growth and development. **D:** Murine C2C12 myoblasts were induced to differentiate in the presence of increasing doses of pharmacological LOX pathway inhibitors including pan LOX inhibitors (ETYA, baicalein, and NDGA), 15-LOX-1 specific inhibitors [BLX-3887, 9c(i472), and ThioLox], platelet type 12-LOX specific inhibitors (ML355 and CAY10698), and 5-LOX specific inhibitors (zileuton, malotilate, or MK886). Myotubes were stained for myosin heavy chain (green) and myogenin (red) following 72 h of myogenic differentiation. Nuclei were counterstained by DAPI (blue). Myotube formation was quantified as percentage of myosin^+^ cell area per field of view. **E:** Primary myoblasts from WT and *Alox15^-/-^*mice were induced to differentiate for 3 days in the presence of a 25 μM dose of various individual long chain (LC) polyunsaturated fatty acids (PUFAs) including n-6 arachidonic acid (ARA), n-3 eicosapentaenoic acid (EPA), n-3 docosapentaenoic acid (DPA), and n-3 docosahexaenoic acid (DHA). Myotubes were stained for myosin heavy chain (green), myogenin (red), and DAPI (blue) for quantitative analysis of myotube diameter. **C-E:** Scale bars are 200 µm. **F-L:** Conditioned media was collected from myotube cultures receiving individual LC-PUFAs including ARA, EPA, DPA, and DHA at a dose of 25 μM or an equimolar mixture of 6.25 μM of each of these individual PUFAs reaching a total concentration of 25 μM. **F:** A heatmap of the top 30 most differentially regulated lipid mediators detected in conditioned culture media samples obtained from primary myotubes derived from WT and *Alox15^-/-^* mice as measured by targeted liquid chromatography-tandem mass spectrometry (LC-MS/MS). **G-L:** Extracellular concentration major representative lipid mediator metabolites of the cyclooxygenase (COX) pathway (e.g., PGE_2_) (**G**), 5-lipoxygenase (5-LOX) pathway (e.g., 7-HDoHE) (**H**), 12-lipoxygenase (12-LOX) pathway (e.g., 14-HDoHE) (**I**), 15-lipoxygenase (15-LOX) pathway (e.g., 17-HDoHE) (**J**), cytochrome P450 (CYP 450) pathway [e.g., 16(17)-EpDPE)] (**K**), and downstream bioactive SPMs (e.g., RvD1) (**L**). Bars show the mean ± SEM of 4-5 mice per group (biological replicates) with dots representing data from each individual mouse. **A-E:** P*-*values were determined by two-tailed unpaired t-tests. *p<0.05 and *P<0.01 vs. WT myotubes**. G-L:** P*-*values were determined by two-way ANOVA followed by Holm-Šídák post-hoc tests. *p<0.05 vs. M0 and #p<0.05 for WT vs. *Alox15^-/-^*cells.

Immunocytochemistry analysis showed that following 72 h of myogenic differentiation that *Alox15^-/-^* myoblasts fused to form smaller myotubes when compared to WT cells (**Fig. 5C**). Treatment of confluent C2C12 myoblasts at the onset of myogenic differentiation with the dual COX/LOX inhibitor eicosatetraynoic acid (ETYA), the pan LOX inhibitor nordihydroguaiaretic acid (NDGA), or the dual 12-LOX/15-LOX inhibitor baicalein each had marked direct suppressive effects on *in vitro* myotube formation (**Fig. 5D, Fig S6A)**. Specific inhibitors of 15-LOX-1 including BLX-3887, 9c(i472), and PD146176 each also dramatically inhibited C2C12 myotube formation at relatively lower doses than pan LOX inhibitors (e.g., 10-20 µM) (**Fig. 5D, Fig S6B)**. Two additional 15-LOX-1 specific inhibitors including ML351 and ThioLox (**Fig S6B**), as well as the platelet-type 12-LOX inhibitor CAY10698 (**Fig S6D**), also dramatically blocked myotube formation, but only at higher doses (100-200 µM) (**Fig. 5D**). Specific inhibitors of the 5-LOX pathway including zileuton and malotilate had little effect on myotube formation at low doses (10-20 µM) (**Fig S6C**). Nevertheless, malotilate (but not zileuton) did markedly blocked myotube formation at higher doses (100-200 µM) (**Fig. 5D)**. Furthermore, MK-866, which interferes with 5-LOX activity indirectly via blockade of 5-LOX activating protein (FLAP) proved to be a highly potent inhibitor of C2C12 myotube formation at relatively low doses (e.g., 25-50 µM) (**Fig. 5D, Fig S6C)**. Overall, these data suggest that leukocyte-type 12/15-LOX (15-LOX-1) and FLAP activity appear to be indispensable for successful muscle cell growth and development *in vitro*.

### LC-PUFA supplementation stimulates SPM biosynthesis and promotes muscle cell growth by a 12/15- LOX dependent pathway

We next studied the potential effect of LC-PUFA supplementation on lipid mediator biosynthesis by isolated skeletal muscle cells. Primary myoblasts obtained from WT and *Alox15*^-/-^ mice were induced to differentiate for 72 h in the presence or absence of a 25 µM dose of pure individual LC-PUFAs including ARA, EPA, DPA, and DHA. Treatment with each of these LC-PUFAs alone greatly increased the mean diameter of WT myotubes (**Fig. 5E**). In contrast, primary myotubes from *Alox15^-/-^* mice failed to respond to supplementation with any of these individual LC-PUFAs (**Fig. 5E**).

LC-MS/MS based profiling detected a total of 100 lipid mediator species present in conditioned culture media obtained from differentiating WT myoblasts in the absence of LC-PUFA supplementation (**Table S1**). Exogenous LC-PUFA treatment markedly increased extracellular concentrations of a wide range of lipid mediators in conditioned media samples (**Fig. 5F**). Supplementation with a 25 μM mixture of ARA, EPA, DPA, and DHA (6.25 μM each) showed the most diverse changes in culture media lipid mediator profile, altering the extracellular concentrations of a total of 40 different species of lipid metabolites (**Table S7**). Additionally, supplementation with a 25 μM dose of pure individual LC-PUFAs including ARA, EPA, DPA, and DHA altered the culture media concentrations of 28, 28, 25, and 32 lipid metabolites, respectively (**Table S7).**

ARA supplementation markedly increased concentrations of many metabolites of the COX pathway (e.g., PGE_2_) (**Fig. 5G, Fig S7A**). Of these, only PGF_2α_ differed between *Alox15*^-/-^ vs WT myotubes with lower levels produced by 12/15-LOX deficient cells (**Fig S7A**). Major 5-, 12-, and 15-LOX metabolites of ARA including 5-HETE (**Fig. S7B**), 12-HETE (**Fig. S7C**) and 15-HETE (**Fig. S7D**) were also greatly increased by ARA supplementation in WT myotubes. In contrast, 5-HETE, 12-HETE and 15-HETE did not increase in *Alox15*^-/-^ cells receiving ARA supplementation (**Fig. S7C-D**). The n-6 PUFA-derived SPM LXA_4_ was also increased following ARA treatment but was not influenced by *Alox15* deficiency (**Fig. S7F**). Nevertheless, the positional lipoxin isomer 15-epi-LXB_4_ was significantly lower in culture media samples obtained from *Alox15*^-/-^ myotubes (**Table. S8**). Supplementation with n-3 EPA greatly increased extracellular concentrations of series-3 COX metabolites (e.g., PGD_3_) (**Table S7**). EPA treatment also greatly increased culture media concentrations of E-series SPMs including 18-HEPE, RvE1, and RvE2 (**Table S7).** Overall, these responses were generally similar in WT and *Alox15*^-/-^ myotubes (**Fig. S7F**, **Table S8)**. Supplementation with n-3 DPA increased concentrations of the di-hydroxy DPA products RvD5_n-3DPA_ and AT-RvD5_n-3DPA_ (**Table S7**), with comparable levels found in WT and *Alox15^-/-^*muscle cells (**Table S8**). Supplementation with n-3 DHA greatly increased concentrations of monohydroxylated 5-, 12-, and 15-LOX metabolites of DHA including 7-HDoHE (**Fig. 5H**), 14-HDoHE (**Fig. 5I**), and 17-HDoHE (**Fig. 5J**). These responses were significantly diminished in *Alox15*^-/-^ muscle cells (**Fig. 5H-J)**. DHA treatment also increased concentrations of some CYP450 metabolites (e.g., 16(17)-EpDPE) in WT, but not *Alox15*^-/-^ muscle cells (**Fig. 5K**). Downstream D-series SPMs including RvD1, RvD2, RvD3, RvD5, PDX, AT-PD1, AT-RvD6, and MaR2 were greatly increased in WT myotubes following DHA supplementation (**Fig. 5L**, **Fig. S7F**). In contrast, *Alox15^-/-^* cultures showed lower extracellular concentrations than WT cells of RvD1, RvD2, RvD5, RvD6, PDX, and AT-PD1 (**Fig. 5L, Fig S7F)**. Similar trends towards diminished levels of PD1, AT-RvD6, and MaR2 were also seen (**Table S8**). Overall, these data suggest that skeletal muscle cells directly produce LOX metabolites *in vitro,* some of which are dependent on the local expression of *Alox15*.

## Discussion

In the current study, we investigated the role of leukocyte-type 12/15-LOX in skeletal muscle inflammation and regeneration in mouse and cell models. *Alox15* knockout dysregulated intramuscular lipid mediator profile leading to basal low-grade inflammation, exaggerated acute immune cell responses to tissue injury, and blunted skeletal muscle regeneration. Mechanistically, 12/15-LOX deficient MФ displayed impaired *in vitro* polarization to a pro-regenerative M2-like phenotype. Furthermore, 12/15-LOX deficient myogenic progenitor cells showed direct defects in *in vitro* myogenesis and were insensitive to the stimulatory effects of LC-PUFA supplementation upon muscle cell growth and differentiation. Overall, these data show that *Alox15* expression is essential for timely inflammation-resolution and regeneration following skeletal muscle injury via multiple mechanisms including both immunomodulation and direct determination of myogenic progenitor cell fate.

*Alox15*^-/-^ mice showed a dysregulated basal intramuscular lipid mediator profile that was associated with chronic low-grade skeletal muscle inflammation, as assessed by intramuscular MФ number. This finding is consistent with prior reports that *Alox15*^-/-^ mice show a chronic inflammatory state in other tissues, such as increased numbers of MФs in the skin^39^ and peritoneal cavity^40^. We hypothesize that basal muscle inflammation in *Alox15*^-/-^ mice could be attributed to a lack of 12/15-LOX derived lipid mediators with anti-inflammatory actions. In this regard, our LC-MS/MS data are generally consistent with a prior report that *Alox15*^-/-^ mice show a basal deficiency of some anti-inflammatory 12/15-LOX metabolites in the skin^39^. Overall, these data suggest that a lack of anti-inflammatory LC-PUFA metabolites may drive chronic low-grade skeletal muscle inflammation in young adult *Alox15*^-/-^ mice similar to our previous observation in aging WT mice^23^.

Chronic low-grade inflammation of skeletal muscle has been implicated in muscle atrophy^41–43^. However, we found little evidence indicative of muscle wasting or weakness in young adult male *Alox15*^-/-^ mice. Nevertheless, we did observe a significant increase in fast-oxidative-glycolytic (type IIA) myofibers in the TA muscle of *Alox15*^-/-^ mice. The potential role of 12/15-LOX activity in muscle fiber type determination remains elusive. When compared to the predominantly fast-twitch TA and EDL muscles, the predominantly slow twitch soleus muscle has been reported to be relatively enriched in 12/15-LOX metabolites^44–46^. Consistently, treatment with RvD2 was found to directly induce satellite cells to differentiate into slow-oxidative (type I) myotubes *in vitro*^44^. In contrast, daily IP injection of RvD2 increased fast-glycolytic fibers (type IIB) fibers in the regenerating TA muscle^44^. The precise role 12/15-LOX in fiber type determination may be muscle-specific, hence, the impact of 12/15-LOX deficiency on the fiber type of different muscles such as the slow-oxidative soleus warrants further study.

We found that *Alox15*^-/-^ mice showed an exaggerated innate immune response to skeletal muscle injury. These data are consistent with prior studies showing that *Alox15*^-/-^ mice mount greater inflammatory responses in several other models (e.g., ^40,47–52^). In contrast, some other experimental models have shown, paradoxically, that *Alox15*^-/-^ mice rather show blunted inflammation (e.g., ^53–59^). 12/15-LOX deficiency differentially impacted a range of inflammatory cytokine/chemokines in the current study. While many classical examples (e.g., TNFα, IL-1β, Il-6, and IL-10) were unaffected, *Alox15*^-/-^ mice showed a far greater local increase in GM-CSF expression following muscle injury. This is consistent with a prior study showing that GM-CSF was one of the most upregulated cytokines in peritoneal MФ isolated from *Alox15*^-/-^ mice^40^. On the other hand, we found the MCP-1 was less robustly induced in *Alox15*^-/-^ mice following muscle injury. This finding is consistent with prior studies showing that 12/15-LOX is an important signal driving MФ expression of MCP-1^60,61^. Notably, MCP-1 plays a crucial role in recruiting blood monocytes to injured skeletal muscle^62,63^. A blunted MCP-1 response in *Alox15*^-/-^ mice has been previously suggested as a potential mechanism to explain reduced monocyte recruitment^61^. Therefore, the increased early MФ infiltration of injured muscle in *Alox15*^-/-^ mice observed in the current study is likely driven by distinct mechanisms independent of MCP-1.

Administration of pharmacological doses of D-series SPMs such as RvD1^23,24^, AT-RvD1^26^, RvD2^22,44,64^, and MaR1^21^ has each recently been shown to limit inflammation, expedite its timely resolution, and stimulate myofiber regeneration following acute skeletal muscle injury. Long term systemic injection of the D-series SPM RvD2 also improved muscle regeneration in the *mdx* mouse model of muscular dystrophy^65,66^. Daily oral gavage with the D-series SPM protectin DX (PDX) was recently reported to be able to protect against development of age-associated musculoskeletal frailty^67^. While most work so far has focused on the n-3 DHA derived D-series SPMs, the E-series resolvin E1 (RvE1) has also been reported to protect muscle cells *in vitro* against inflammation and atrophy induced by lipopolysaccharide (LPS)^68^. Based on these prior studies we hypothesized that the heightened acute immune response following muscle injury in *Alox15*^-/-^ mice might be associated with a lack of SPMs and/or their pathway markers. Consistent with our prior studies^23,24^, we observed clear increases in local concentrations of 14-HDoHE and 17-HDoHE following muscle injury in WT mice. We also detected increased concentrations of some downstream DHA-derived SPMs including RvD1, RvD6, AT-RvD3, and MaR1. Surprisingly, absolute intramuscular concentrations of these analytes were clearly not lacking in *Alox15*^-/-^ mice. Nevertheless, the ratio of DHA derived SPMs and their pathway markers (e.g., 14-HDoHE and 17-HDoHE) relative to pro-inflammatory eicosanoids (e.g., PGE_2_) was reduced in 12/15-LOX deficient mice. As such, a relative deficiency of pro-resolving lipid mediators in combination with an absolute overabundance of pro-inflammatory eicosanoids may still contribute to the observed exaggerated acute immune cell responses in the current study.

Most of the studies conducted so far have focused on the impact of D-series SPMs on skeletal muscle injury and repair. Nevertheless, intravenous injection of the n-6 ARA-derived SPM LXA_4_ was also found to limit skeletal muscle inflammation following ischemia reperfusion injury in rats^69^. Consistent with our prior studies^23,24^, both LXA_4_ and its precursor molecule 15-HETE were greatly elevated after muscle injury in the current study. Surprisingly, 15-HETE and LXA_4_ were both present at greater concentrations in *Alox15*^-/-^ mice when compared to WT controls. This might be explained, in part, by the existence of alternative 15-LOX-1 independent routes of lipoxin biosynthesis. For example, cells expressing 5-LOX can convert ARA to leukotriene A_4_ (LTA_4_) which can be taken up by platelet-type 12-LOX expressing cells and converted to LXA_4_ and LXB_4_^70^. We found that both 5-LOX (*Alox5*) and platelet-type 12-LOX (*Alox12*) were more highly expressed in *Alox15*^-/-^ mice following muscle injury. Thus, the 5-LOX/12-LOX pathway may contribute to lipoxin biosynthesis in injured muscle. Prior studies have further shown that, substantial quantities of 15-HETE can be formed via COX-1 activity in 12/15-LOX knockout MФs, especially when 5-LOX was inactivated in parallel^71^. Indeed, herein COX-1 was more highly expressed in *Alox15*^-/-^ mice following muscle injury. Therefore, the COX-1 pathway might be an additional important contributor to 15-HETE and/or LXA_4_ biosynthesis in injured skeletal muscle.

We observed that BMMs obtained from *Alox15*^-/-^ mice exhibited exaggerated M1 polarization as well as impaired M2 polarization, and this was associated with underlying defects in lipid mediator class switching. These findings are consistent with prior studies reporting that certain 12/15-LOX metabolites, such as RvD1, can induce M2 MФ polarization^72^. Additionally, these findings agree with prior reports that M2 MФs may produce relatively greater concentrations of 12/15-LOX pathway metabolites when compared to M1 MФ^73,74^. While it was originally suggested that peritoneal MФ obtained from *Alox15*^-/-^ mice were indistinguishable from WT^30^, subsequent research showed that *Alox15*^-/-^ peritoneal MФ are greatly impaired in their ability to phagocytose apoptotic cells (efferocytosis)^48,75,76^. Moreover, consistent with our data, 12/15-LOX deficient peritoneal MФ were previously reported to be deficient in induction of the scavenger receptor CD36 in response to treatment with the anti-inflammatory cytokine IL-4^77^. Overall, these data suggest that impaired MФ functions may contribute, in part, to delayed inflammation-resolution in 12/15-LOX knockout mice.

We also found that *Alox15*^-/-^ mice displayed impaired *in vivo* myofiber regeneration. This finding is consistent with prior reports that *Alox15*^-/-^ mice display poor epithelial wound healing^78,79^. In contrast, *Alox15*^-/-^ mice have rather been found to be protected against inflammation-induced cardiac muscle damage^53–56,80^. The apparent overall deleterious effects of 12/15-LOX deficiency on skeletal muscle regeneration in the current study may be related to dysregulated immune-muscle cell interactions. PMNs can inflict secondary myofiber injury and delay myofiber regeneration following muscle injury^81^. Therefore, an excessive intramuscular PMN presence and impaired clearance from the site of injury in *Alox15*^-/-^ mice may have exacerbated local tissue damage and impeded myofiber regeneration. Blood monocytes are known to play an important supportive role in skeletal muscle regeneration, in part, by engulfing and clearing necrotic myofiber debris^82^. Nevertheless, earlier than normal peak blood monocyte/M1 MФ infiltration in *Alox15*^-/-^ mice may have negatively impact tissue repair, especially if their phagocytic capacity is impaired^48,75,76^. Surprisingly given that *Alox15*^-/-^ BMMs display defective M2 polarization *in vitro*, we observed greater numbers of M2-like CD163^+^ MФ within regenerating muscles of *Alox15*^-/-^ mice. M2 macrophages have generally been thought to play a beneficial role in skeletal muscle regeneration^83–85^. Nevertheless, it has also been reported that *Cd163*^-/-^ mice display improved skeletal muscle regeneration in a model of limb ischemia^86^. Another recent study suggested that depletion of CD206^+^ M2-like MФ also enhanced muscle regeneration following cardiotoxin induced injury^87^. Therefore, it is possible that an earlier than normal local transition of the greater numbers of infiltrating blood monocytes to M2 MФ in *Alox15*^-/-^ mice could interfere with muscle regeneration via multiple mechanisms^86,87^.

In addition to evidence of dysregulated immune-muscle cell crosstalk in 12/15-LOX deficient mice, we found that myogenic progenitor cells isolated from muscle tissue of *Alox15*^-/-^ mice showed marked impairments in their ability to form mature myotubes when cultured *in vitro*. Pharmacological LOX inhibitors also proved to be robust inhibitors of myotube formation in WT muscle cells. Amongst the pan LOX inhibitors tested, NDGA was the most potent inhibitor of myogenesis in our hands. This finding is consistent with a previously published paper showing that NDGA treatment resulted in severe dose-dependent inhibition of C2C12 myoblast differentiation^88^. Interestingly, the authors concluded that this marked inhibitory effect on *in vitro* myogenesis was independent of LOX since selective inhibitors of 5-LOX and platelet-type 12-LOX did not mimic the suppressive effects of NDGA^88^. Nevertheless, selective inhibitors of 15-LOX-1 were not tested^88^. When combined with our results, it appears likely that the deleterious effects of NDGA on myotube formation is primarily attributable to inhibition of 15-LOX-1. We found that the FLAP inhibitor MK886 also blocked C2C12 myotube formation, suggesting that FLAP activity is also indispensable for myotube formation. Interestingly, leukocyte FLAP activity has been previously shown to be required for the sequential 5-LOX mediated oxygenation of 15-HETE and/or 17-HDoHE to form downstream di- and tri-hydroxylated SPMs^89–91^.

It has been previously suggested that 15-HETE, the primary 15-LOX metabolite of n-6 ARA, is a cachexic factor produced by skeletal muscle cells that promotes protein degradation^92–95^. Nevertheless, consistent with prior published work by us and others^33,96–100^, ARA supplementation greatly stimulated *in vitro* myogenesis of WT myoblasts in the current study. In contrast, *Alox15*^-/-^ myoblasts were unresponsive to the stimulatory effects of ARA on myotube formation, and this was associated with lower 15-HETE levels. Consistent with some prior studies, we found that supplementation with n-3 LC-PUFAs including EPA, DPA, and DHA each also markedly stimulated *in vitro* myotube formation^97,101–106^. Like ARA, the stimulatory actions of n-3 LC-PUFAs on *in vitro* myogenesis were also dependent on 12/15-LOX activity. A lack of muscle cell growth in *Alox15*^-/-^ myoblasts in response to n-3 DHA supplementation was associated with diminished extracellular concentrations of many 12/15-LOX metabolites of DHA including several D-series SPMs. Overall, these data are consistent with recent reports by us and others that RvD1^24^, RvD2^44,65,66^, RvD3^107^, MaR1^21^, and RvE1^68^ have direct pro-myogenic and/or anti-catabolic effects on skeletal muscle cells cultured *in vitro*.

Taken together, our data strongly demonstrate that leukocyte-type 12/15-LOX is an important determinant of timely resolution of inflammation and myofiber regeneration following skeletal muscle injury. Consistent with previously published work, our data suggest that *Alox15* is important for MФ polarization to pro-resolutive and tissue reparative phenotype. Finally, we report a novel and direct role of *Alox15* as a previously unknown important intrinsic determinant of myogenic progenitor cell fate.

## Study approval

All animal experiments were approved by the University of Michigan (PRO00008744) and Purdue University (2205002271) Institutional Animal Care and Use committees (IACUC).

## Author contributions

J.F.M conceived the study. J.F.M, S.V.B, H.V.R, K.R.M, and S.K supervised the work. J.F.M, B.S, X.L, H.R, V.C.F, and S.K designed the experiments. J.F.M, C.D, B.S, H.R, V.C.F, and X.L performed the experiments. J.F.M, B.S, X.L, H.R, B.S, C.D, and K.R.M analyzed the data. J.F.M, B.S, H.R, and XL prepared the figures and wrote the manuscript with input from all authors

## Acknowledgments

This work was supported, in part, by grants from the National Institutes of Health (NIH) including S10RR027926 (KRM), S10OD032292 (KRM), R01DK132819 (SK), and R01AG050676 (SVB), together with the Purdue University College of Agriculture (JFM), Purdue University Department of Animal Sciences (JFM), and the Ralph W. and Grace M. Showalter Research Trust (JFM). Monoclonal antibodies including F1.652s, BA-D5c, SC-71c, BF-F3c, MF-20, and F5D were obtained from the Developmental Studies Hybridoma Bank (DSHB), created by the NICHD of the NIH and maintained at The University of Iowa, Department of Biology, Iowa City, IA 52242.

---

**Supplementary Figure 1:**
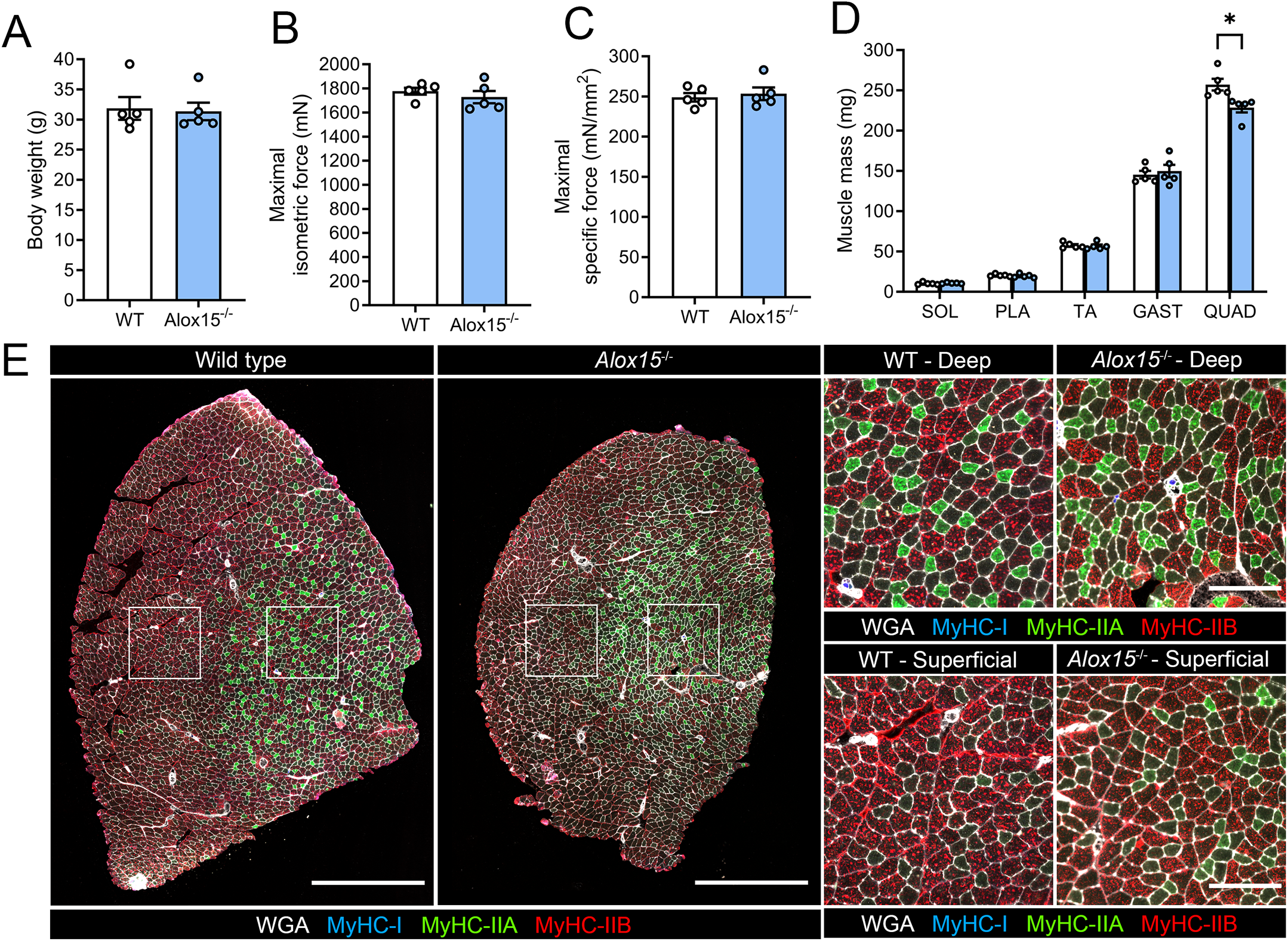
Characterization of the basal skeletal muscle phenotype of 12/15-LOX deficient mice. **A:** Body weights (g) of wild type (WT) and *Alox15* knockout (*Alox15*^-/-^) mice. **B:** Measurement of *in situ* absolute maximal isometric force (mN) produced by the tibialis anterior (TA) muscle of WT and *Alox15*^-/-^ mice. **C:** Maximal specific isometric contractile force (sP_O_), mN/mm²) of WT and *Alox15*^-/-^ mice. **D:** Absolute muscle mass of soleus (SOL), plantaris (PLA), tibialis anterior (TA), gastrocnemius (GAST), and quadriceps (QUAD) muscles from WT and *Alox15*^-/-^ mice. **E:** TA cross-sections from WT and *Alox15*^-/-^ mice were stained with antibodies against myosin heavy chain (MyHC) isoforms I, IIA, and IIB to identify specific muscle fiber types. Type IIx fibers remain unstained (black). Bars show the mean ± SEM of 4-5 mice per group (biological replicates) with dots representing data from each individual mouse. P*-*values were determined by two-tailed unpaired t-tests. *p<0.05 WT vs. *Alox15*^-/-^ mice. Scale bars are 1000 µm for stitched whole muscle sections and 200 µm for representative fields of view.

**Supplementary Figure 2:**
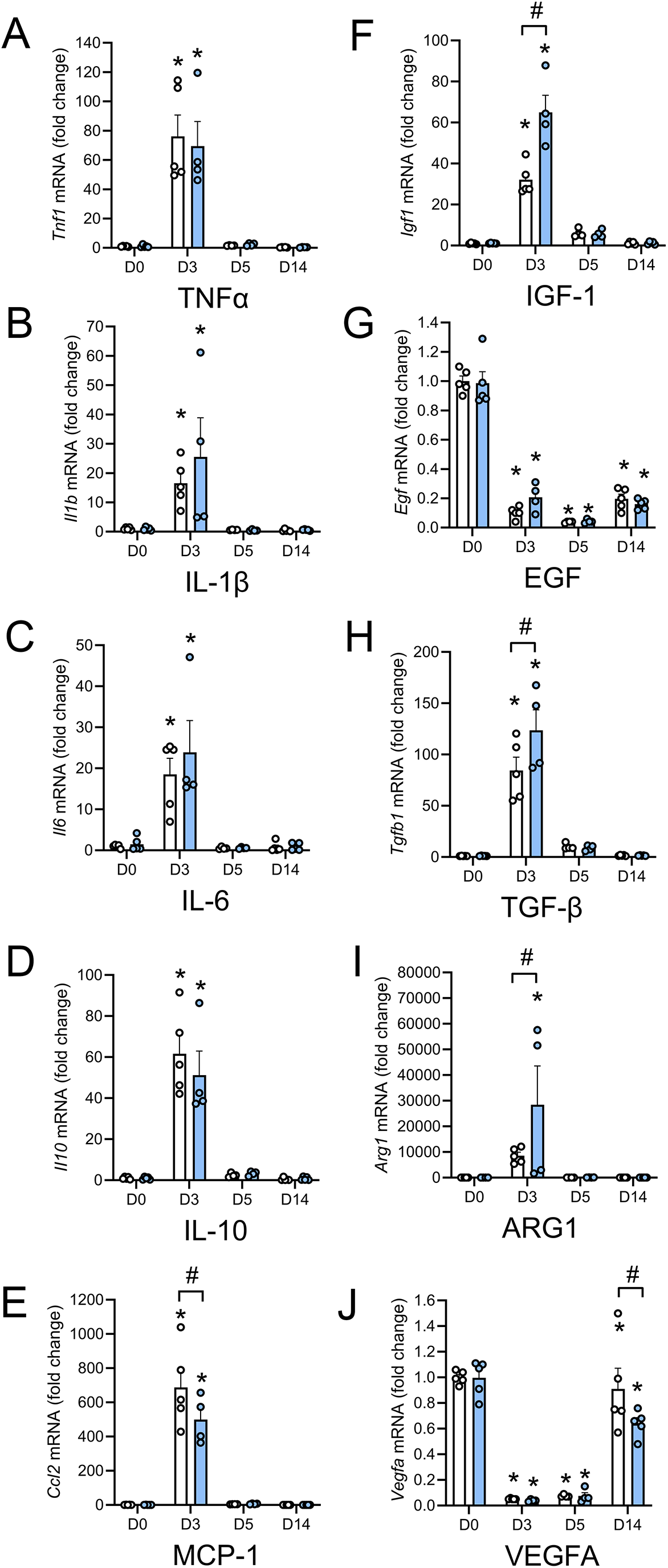
Effect of 12/15-LOX knockout on local expression of inflammatory cytokines and growth factors following skeletal muscle injury. **A-E:** Muscle mRNA expression of cytokines/chemokines including TNFα (*Tnf1*) (**A**), IL-1β (*Il1b*) (**B**) IL-6 (*Il6*) (**C**), IL-10 (*Il10*) (**D**), and MCP-1 (*Ccl2*) (**E**) at day 3 (D3), day 5 (D5), and day 14 (D14) following muscle injury induced by intramuscular injection of barium chloride (BaCL_2_). **F-J:** Muscle mRNA expression of growth factors including IGF-1 (*Igf1*) (**F**), EGF (*Egf*) (**G**), TGF-β (*Tgfb1*) (**H**), arginase 1 (*Arg1*) (**I**), and VEGF (*Vegfa*) (**J**) at D3, D5, and D14 following muscle injury induced by intramuscular injection of BaCL_2_. Bars show the mean ± SEM of 4-5 mice per group (biological replicates with dots representing data from each individual mouse. P*-*values were determined by two-way ANOVA followed by Holm-Šídák post-hoc tests. *p<0.05 vs. D0 and #p<0.05 for WT vs. *Alox15^-/-^*mice.

**Supplementary Figure 3:**
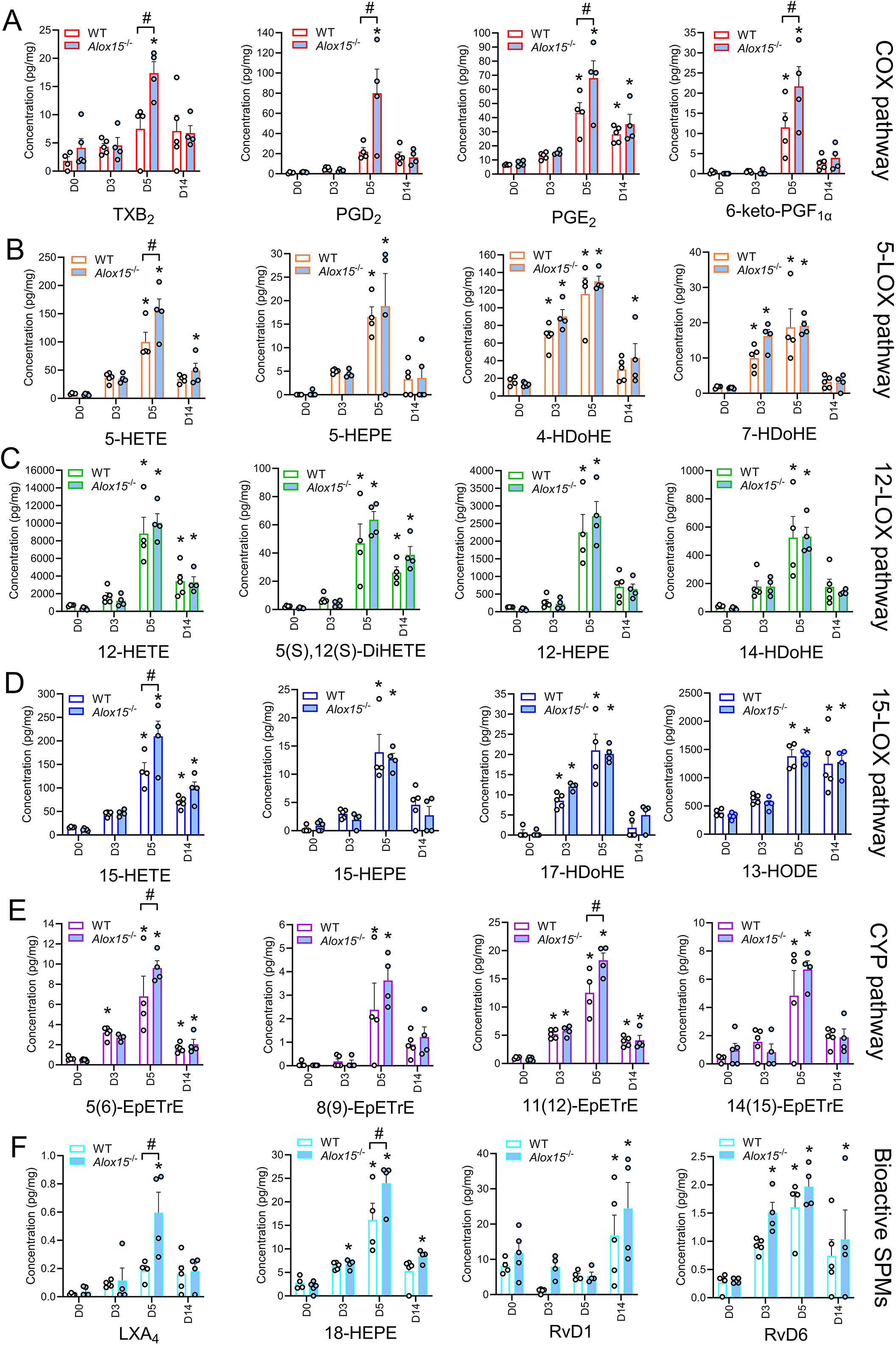
Metabolipidomic profiling reveals a local imbalance of pro-inflammatory vs. anti-inflammatory/pro-resolving lipid mediators following muscle injury. **A-F:** Intramuscular concentrations of major metabolites of the COX pathway (**A**), 5-LOX pathway (**B**), 12-LOX pathway (**C**), 15-LOX pathway (**D**), CYP pathway (**E**), and downstream bioactive SPMs (**F**) detected at day 3 (D3), day 5 (D5), and day 14 (D14) following muscle injury induced by intramuscular injection of BaCl_2_. Bars show the mean ± SEM of 4-5 mice per group (biological replicates) with dots representing data from each individual mouse. P*-*values were determined by two-way ANOVA followed by Holm-Šídák post-hoc tests. *p<0.05 vs. D0 and #p<0.05 for WT vs. *Alox15^-/-^*mice.

**Supplementary Figure 4:**
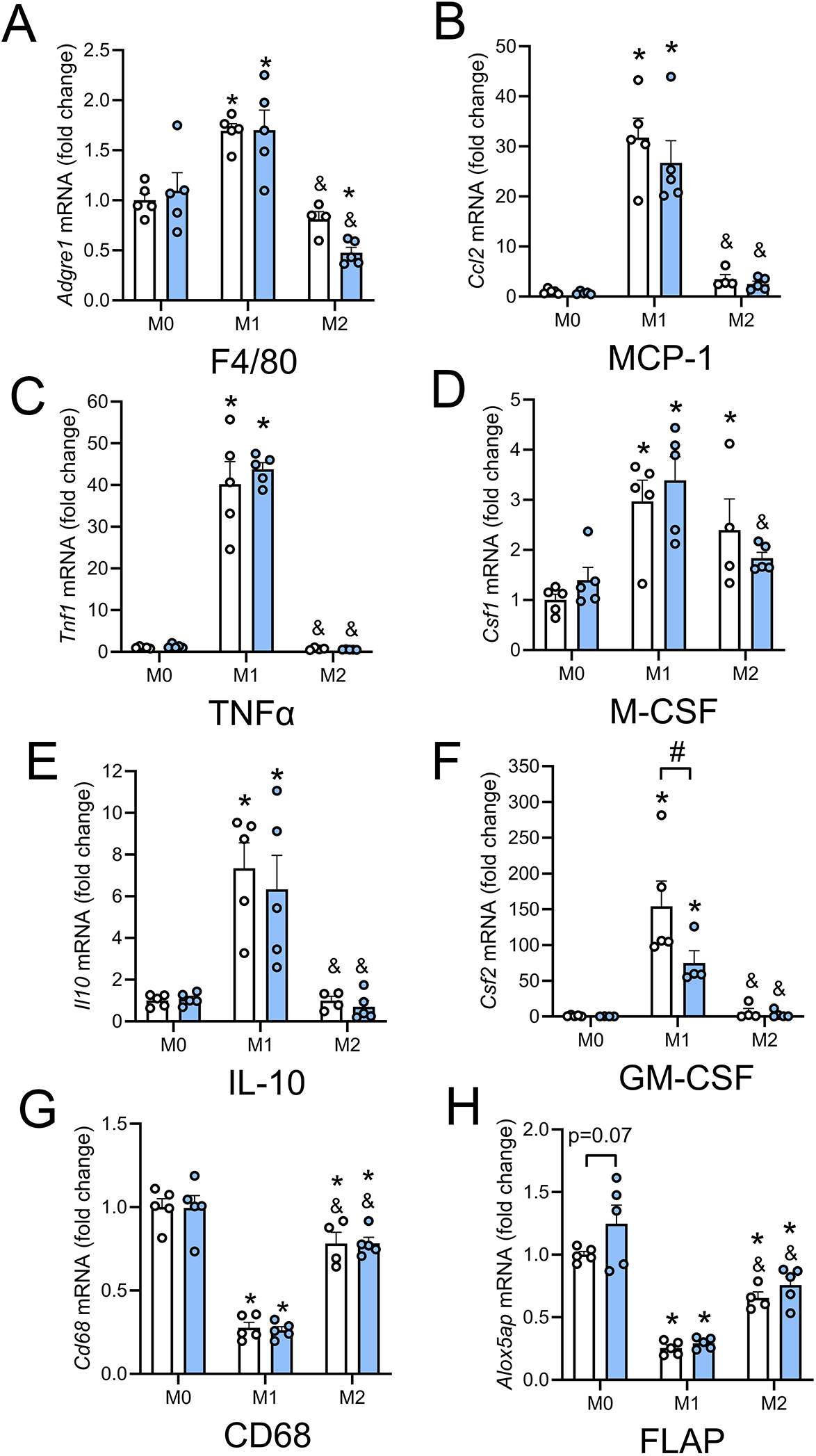
Effect of 12/15-LOX knockout on expression of inflammation related genes in bone marrow derived macrophages. Bone marrow cells were cultured for 7 days in the present of M-CSF to obtain adherent bone marrow-derived macrophages (BMMs). BMMs were maintained in serum free media lacking M-CSF for 24 h to obtain naïve M0 BMM, or polarized to a M1 or M2 phenotype by 24 h treatment with LPS (100 ng/mL) + INF-γ (20 ng/mL) or IL-4 (20 ng/mL), respectively. Expression of mRNA encoding F4/80 (*Adgre1*) (**A**), MCP-1 (*Ccl2*) (**B**), TNFα (*Tnf*) (**C**), M-CSF (*Csf1*) (**D**), IL-10 (*Il10*) (**E**), GM-CSF (*Csf2*) (**F**), CD68 (*Cd68*) (**G**), and FLAP (*Alox5ap*) (**H**) in BMMs obtained from wild type (WT) and 12/15-LOX deficient (*Alox15*^-/-^) mice. Gene expression was measured via RT-qPCR. P*-*values were determined by two-way ANOVA with Holm-Šídák post-hoc tests. *p<0.05 vs. M0, &p<0.05 vs. M1, and #p<0.05 for WT vs. *Alox15^-/-^*cells.

**Supplementary Figure 5:**
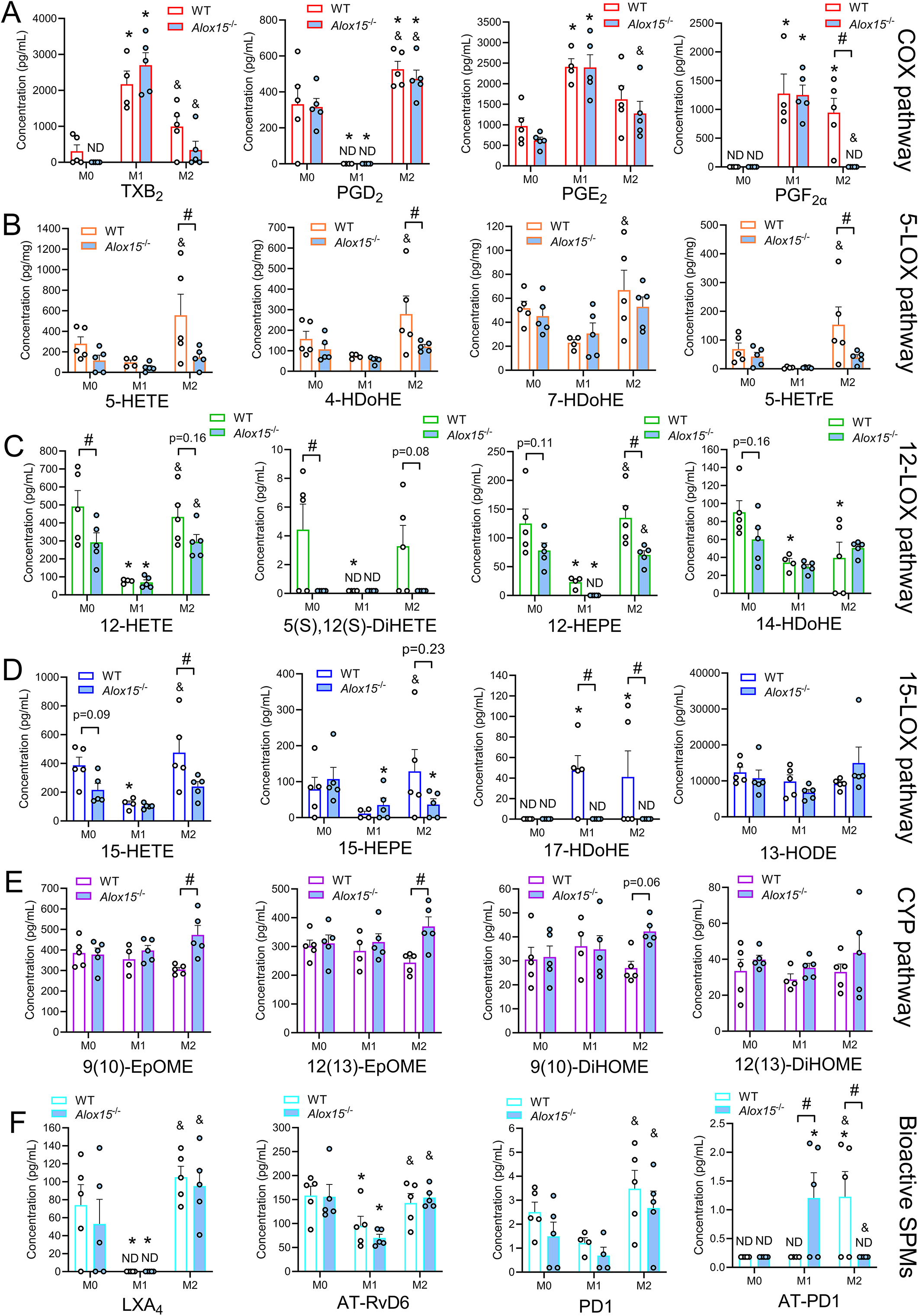
Bone marrow-derived macrophages from *Alox15*^-/-^ mice are deficient in 12/15-LOX-derived lipid mediators under M2-polarizing conditions. **A-F:** Extracellular concentrations of major metabolites of the cyclooxygenase (COX) pathway (**A**), 5-lipoxygenase (5-LOX) pathway (**B**), 12-lipoxygenase (12-LOX) pathway (**C**), 15-lipoxygenase (15-LOX) pathway (**D**), cytochrome p450 (CYP) pathway (**E**), and downstream bioactive specialized pro-resolving mediators (SPMs) (**F**) in serum free conditioned cell culture media samples obtained from M0, M1, or M2 bone-marrow-derived macrophage (BMM) cultures from wild type (WT) and 12/15-LOX deficient (*Alox15*^-/-^) mice. Bars show the mean ± SEM of BMMs obtained from n=5 mice per group (biological replicates) with dots representing BMMs from each individual mouse. P*-*values were determined by two-way ANOVA with Holm-Šídák post-hoc tests. *p<0.05 vs. M0, &p<0.05 vs. M1, and #p<0.05 for WT vs. *Alox15^-/-^*cells.

**Supplementary Figure 6:**
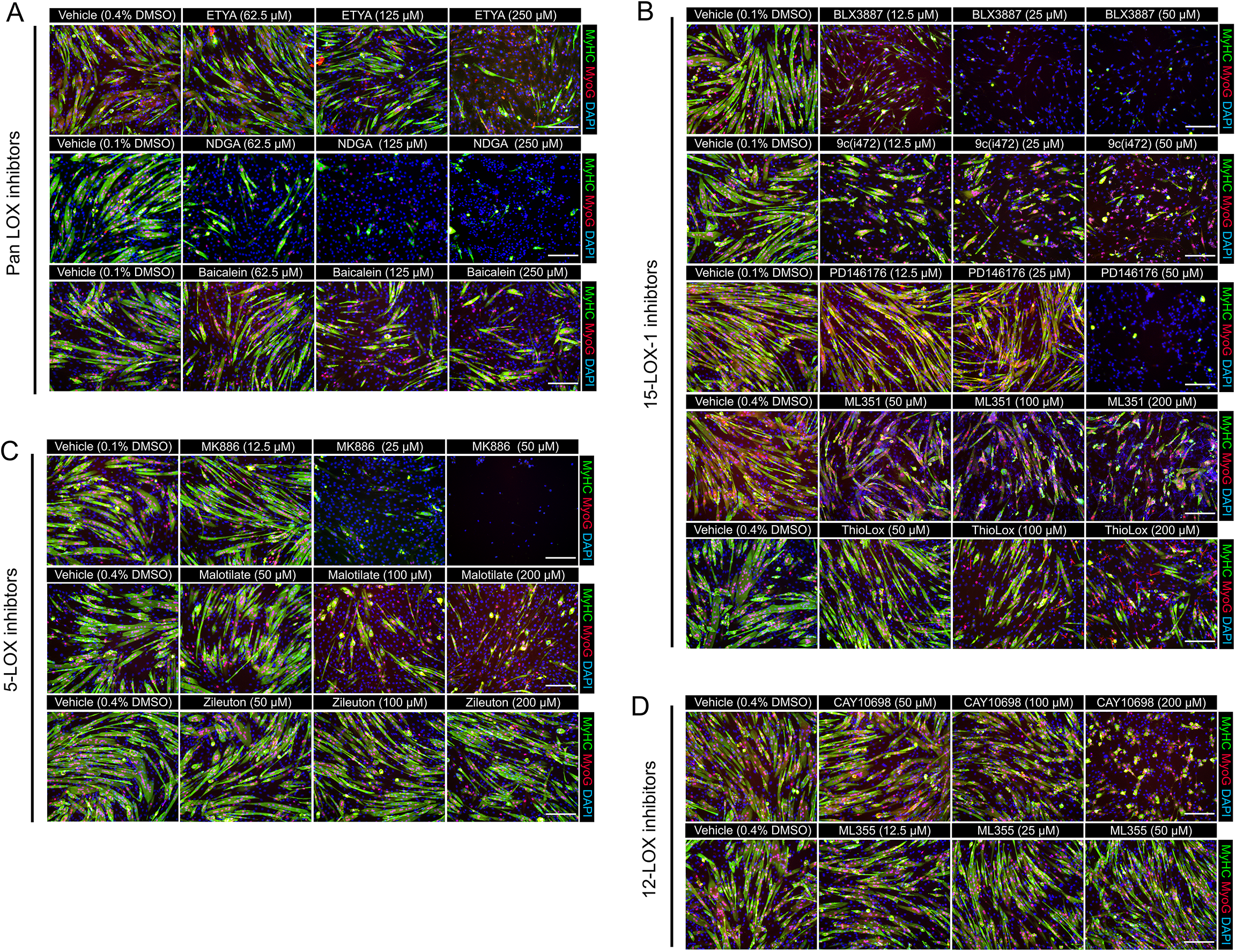
Pharmacological inhibition of 15-LOX-1 and FLAP markedly suppresses *in vitro* myogenesis. Murine C2C12 myoblasts were grown to confluence and then induced to undergo myogenic differentiation in the presence of various doses of pharmacological LOX enzyme inhibitors. (**A**): Pan LOX inhibitors including eicosatetraynoic acid (ETYA), nordihydroguaiaretic acid (NDGA), and baicalein. (**B**): 5-LOX specific inhibitors including MK886, malotilate, and zileuton. (**C**): 12-LOX specific inhibitors including CAY10698 and ML355. (**D**): 15-LOX-1 specific inhibitors including BLX3887, 9c(i472), PD146176, ML351, and ThioLox. Following 72 h of myogenic differentiation resulting myotube cultures were fixed in 4% PFA and stained with primary antibodies against sarcomeric myosin heavy chain (MyHC, green), myogenin (MyoG, red). Cell nuclei were counterstained with DAPI (blue). Results are representative of 3 independent experiments. Scale bars are 200 µm.

**Supplementary Figure 7:**
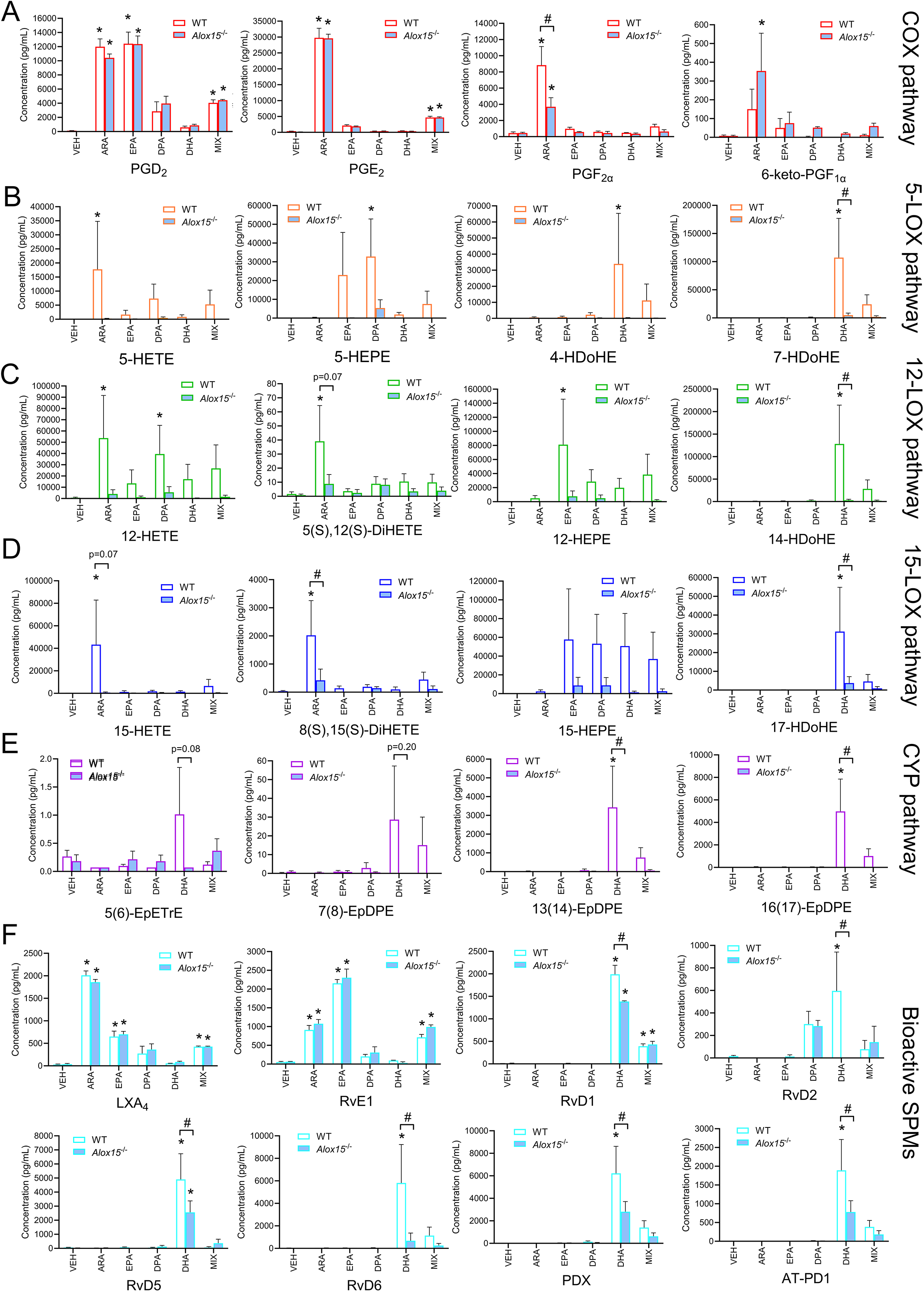
Myogenic progenitor cells produce specialized pro-resolving mediators (SPMs) via a 12/15-LOX dependent pathway. Primary mouse myoblasts were induced to differentiate for 72 h in culture media supplemented with vehicle control (0.02% ethanol), a 25 µM dose of individual pure long chain polyunsaturated fatty acids (LC-PUFAs) including arachidonic acid (ARA), eicosapentaenoic acid (EPA), docosapentaenoic acid (DPA), docosahexaenoic acid (DHA), or an equimolar mixture of ARA, EPA, DPA, and DHA (6.25 µM each). Extracellular concentrations of major metabolites of the COX pathway (**A**), 5-LOX pathway (**B**), 12-LOX pathway (**C**), 15-LOX pathway (**D**), CYP pathway (**E**), and downstream bioactive SPMs (**F**) as detected by LC-MS/MS analysis of conditioned culture media samples obtained from differentiating wild type (WT) and *Alox15*^-/-^ myoblasts. Bars show the mean ± SEM of primary myoblasts obtained from n=4 mice per group. P*-*values were determined by two-way ANOVA followed by Holm-Šídák post-hoc tests. *p<0.05 vs. vehicle and #p<0.05 for WT vs. *Alox15^-/-^* myotubes.

